# Frequent allopolyploidy with distant progenitors in the moss genera *Physcomitrium* and *Entosthodon* (Funariaceae) identified via subgenome phasing of targeted nuclear genes

**DOI:** 10.1101/2023.07.17.549320

**Authors:** Nikisha Patel, Rafael Medina, Lindsay D. Williams, Olivia Lemieux, Bernard Goffinet, Matthew G. Johnson

## Abstract

Allopolyploids represent a new frontier in species discovery among embryophytes. Within mosses, allopolyploid discovery is challenged by low morphological complexity. The rapid expansion of sub-genome sequencing approaches in addition to computational developments to identifying genome merger and whole-genome duplication using variation among nuclear loci representing homeologs has allowed for increased allopolyploid discovery among mosses. We confirm the intergeneric hybrid nature of *Entosthodon hungaricus*, and the allopolyploid origin of *P. eurystomum* and of one population of *P. collenchymatum*. We also reveal that hybridization gave rise to *P. immersum*, as well as to yet unrecognized lineages sharing the phenotype of *P. pyriforme*, and *P. sphaericum.* Our findings demonstrate the utility of a novel approach to phasing homeologs within loci and phasing loci across subgenomes, or subgenome assignment, called homologizer, when working with polyploid genomes, and its value in identifying progenitor species using target capture data.

## Introduction

The accurate quantification of species diversity is central to evolutionary biology. To this end, a molecular phylogenetic approach is indispensable, though it is increasingly recognized that, in plants, lineages of hybrid origin are frequently excluded from such analyses (Soltis et al., 2007; Barker et al., 2016; Patel et al., 2021). Allopolyploidy is a form of polyploidy in which a genome comprises multiple subgenomes each inherited from two or more distinct progenitor species, and such merger may serve as potential mechanism of speciation (Soltis et al., 2015; Lohaus & van der Peer, 2016). The accurate identification of allopolyploids and their inclusion in phylogenetic analyses is, however, impeded by their tendency to be morphologically cryptic, as well as by challenges in phasing homeologous sequences within loci and accurately assigning the variants of all loci to parental subgenomes (Rothfels, 2021). The phylogenetic placement of allopolyploid subgenomes is critical for the identification of parental lineages and therefore a fuller understanding of trait evolution, ecology, and biogeography of groups including allopolyploids and their subgenomes (Oxelman et al., 2017; Rothfels et al., 2017; Rothfels, 2021). The use of DNA data in the identification of allopolyploid parents has been widely accomplished in crop plants including *Gossypium* and *Triticum* (Ozkan et al., 2001; Grover et al., 2015; Chen et al., 2020), but methodological challenges remain in accomplishing this on a large scale for non-model plant species.

With a molecular approach, allopolyploid detection relies on the identification of variation within loci representing homeologs. We refer to homeologs, the copies from each of the progenitors of an allopolyploid and which are assumed to have no recombination, in contrast to allelic variation that segregates at a locus. In addition, here, heterozygosity refers to SNPs distinguishing homeologs from each subgenome in an allopolyploid genome. Historically, studies of polyploid species complexes have detected allopolyploids and identified homeologs by assessing nuclear markers for evidence of heterozygosity in the form of mixed nucleotide signals (for Sanger sequencing) or multiple allele sizes (for microsatellites; Tate et al., 2006; Jorgensen & Barrington, 2017; Clark et al., 2017; Patel et al., 2018).

Phylogenetic inference is also used in identifying allopolyploid lineages. Historically, incongruence in phylogenetic position between nuclear and plastid trees was the basis to identify potential hybrids. However, an organellar genome represents only the maternal heritage is considered a single gene by virtue of lacking recombination. Hence, phylogenetic inference increasingly relies on nuclear genes (Doyle, 2021). Although nuclear genetic markers are biparentally inherited, if phylogenetic inference is based on them without regard for within-sample variation (i.e., assembling reads to produce consensus sequences) sequences could be chimeric and not correctly resolve allopolyploid lineages (Oxelman et al., 2017; Šlenker et al., 2021). Single molecule methods representing individual alleles or homeologs generated via vector cloning or 2-step PCR have been used to manually identify homeologs for a few discrete nuclear genes (Tate et al. 2006; Beike et al. 2014; Jorgensen et al. 2017), but such an approach is not applicable to larger sets of loci, which may be needed to enhance accuracy in parental identification, particularly in cases of hybridization following rapid radiation or incomplete lineage sorting.

High-throughput sequencing methods have substantially improved phylogenetic resolution for non-polyploid species, but inclusion of polyploids in phylogenetic analysis has been more challenging. Genome reduction approaches such as Hyb-Seq have allowed for rapid sequencing of hundreds of low-copy nuclear genes (Weitemier et al., 2014; Hale et al., 2020), and species trees are typically estimated under the coalescent model from unrooted gene trees. This approach provides data for robust phylogenetic inference and, by using pipelines such as HybPiper (Johnson et al., 2016) and HybPhaser (Nauheimer et al., 2021), allows for the discovery of likely allopolyploids through the detection of multiple copies and high heterozygosity (SNPs distinguishing homeologs) within biparentally inherited nuclear markers. However, beyond the detection of allopolyploids, the identification of progenitor lineages is crucial to elucidating evolutionary history. Although Hyb-Seq data show promise in detection of allopolyploids through identification of progenitors based on the phylogenetic affinity of homeologs, significant challenges remain in subgenome phasing, e.g. the correct assignment of each homeologs to parental subgenomes. Some studies have used parallel amplicon sequencing to generate a few nuclear loci and identified homeologs via manual comparison of gene trees. These approaches avoid problems associated both with read assembly by utilizing long-reads sequencing such as PacBio, but scaling this analytical approach to the number of loci typical to a target capture approach like HybSeqis not feasible (Rothfels et al., 2017).

To incorporate sequences representing subgenomes in a phylogeny, the homeolog sequences must be *in phase*, meaning that homeologs are assigned to the correct subgenome across loci. Within loci, tools such as ReadBackedPhasing in the Genome Analysis Toolkit can associate neighboring sequence variants using the sequence reads within a locus (Francioli et al., 2017). Across loci, additional evidence is needed to assign single-locus phased homeologs to each subgenome. Recently developed computational approaches allow for the subgenome phasing of dozens to hundreds of loci and hence the identification of hybrid progenitors. HybPhaser (Nauheimer et al., 2021) uses a two-step phylogenetic approach that identifies putative hybrids and re-assembles sequences using clade affinities, therefore requiring *a priori* identification of appropriate references for each subgenome. The AlleleSorting pipeline (Šlenker et al., 2021) uses a distance-based approach to separate homeologs and assign them to subgenomes. Here, we infer the subgenome phase of sequences using the Homologizer method (Freyman et al., 2023), implemented in RevBayes (Höhna et al., 2016). Homologizer differs from the other recently developed methods by inferring the subgenome phase simultaneously with a phylogeny that includes the phased subgenomes of any allopolyploids as separate tips. Accordingly, an *a priori* reference sequence is not required.

Among mosses, shifts in ploidy arise from allopolyploidy or genome merger (Natcheva & Cronberg, 2004; Swangprogh & Cronberg, 2021) or from autopolyploidy, that is intraspecific genome doubling (Patel et al., 2021). Historically, allopolyploids have been considered both rare and insignificant to moss evolution (Vitt, 1971; Smith, 1978). However, molecular phylogenetic studies in lineages such as Sphagnaceae (Ricca & Shaw, 2010), Mniaceae (Wyatt et al., 1988), and Funariaceae (McDaniel et al., 2010; Beike et al., 2014) suggest that allopolyploids may figure prominently in the evolution of mosses. Allopolyploid mosses may, however, not be morphologically distinct from, or intermediate to, their progenitors, and the identification of the hybrid origin of taxa, and hence of significant morphologically cryptic diversity in mosses, would thus rely on exploring molecular data for extensive fixed heterozygosity indicative of allopolyploid genomes. Accordingly, the exploration of molecular data toward the discovery of novel polyploid lineages has the potential to reveal significant morphologically cryptic diversity in mosses.

Our implementation of the homologizer method to discover polyploids and their progenitors focuses on the Funariaceae, a cosmopolitan family of approximately 300 annual moss species (Crosby et al., 2000; McIntosh, 2007). Most species in this family reside in the genera *Funaria, Entosthodon,* and *Physcomitrium* (Fife, 1985). The Funariaceae, like several other moss families, include several complexes of putative, and likely unrecognized, polyploid species (Patel et al., 2021; note that when speaking of the ploidy level of mosses, it is customary to use the gametophyte generation as the standard — thus a “diploid” moss is a polyploid and it has a tetraploid sporophyte). Like other bryophytes, the life cycle of Funariaceae comprises a free-living vegetative haploid generation, the gametophyte, with, following sexual reproduction, a permanently attached diploid generation, the sporophyte (Figure 1). The Funariaceae exhibit narrow phenotypic diversity in the vegetative haploid gametophyte in contrast to extensive trait variation in the diploid sporophyte such as seta length, peristome architecture, or sporangial complexity and symmetry (Fife, 1985).

**Figure 1:**
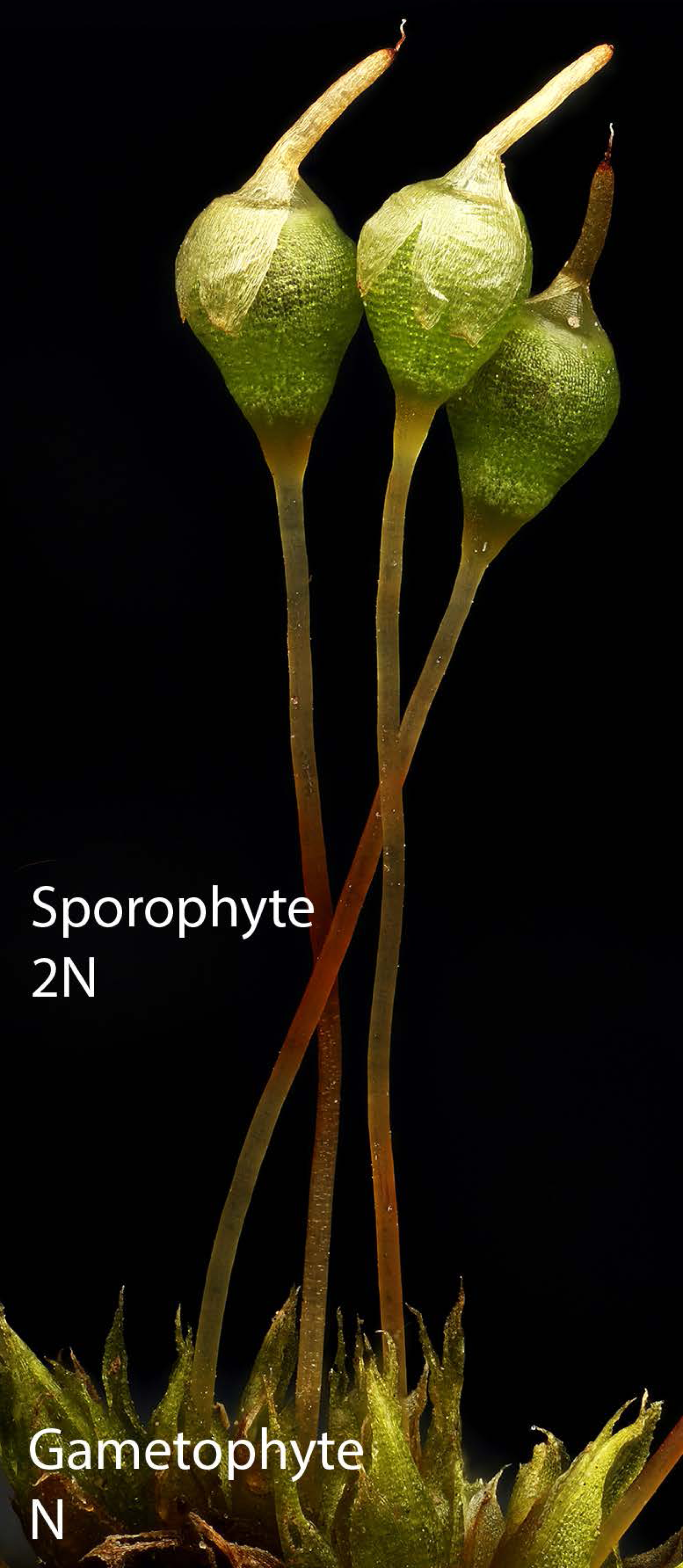
The gametophytic (haploid 1N) and sporophytic (diploid 2N) generations of *Physcomitrium pyriforme s. lato*.

In this study we aim **(1)** first to confirm the allopolyploidy of three species of the Funariaceae using 50 nuclear genes: *Entosthodon hungaricus, Physcomitrium, eurystomum,,* and *P. collenchymatum,* and investigate new evidence of allopolyploidy in species, *P. immersum, P. sphaericum,* and *P. pyriforme.* All species except *P. collenchymatum* were sampled for target capture sequencing in Medina et al. (2019) but excluded from phylogenetic analysis as they showed heterozygosity congruent with a potential allopolyploid origin. The ploidy and evolutionary origins of *P. collenchymatum* are additionally considered here to address previously conflicting reports of allopolypoidy associated with the *P. collenchymatum* phenotype. McDaniel et al. (2010), Beike et al. (2014), and Ostendorf et al. (2021), each using the same nuclear dataset, report, based on a single identical population, that *P. collenchymatum* is an allopolyploid whereas Medina et al. (2018; 2019) treated other populations with this phenotype as haploid. Our second aim **(2)** is to infer evolutionary origins for component subgenomes and in doing so **(3)** highlight the utility and drawbacks of Homologizer as an approach to subgenome phasing and in identifying maternal and paternal progenitors of hybrid species. The development and implementation of a new approach to the discovery and characterization of these lineages will allow us to better understand the contribution of allopolyploidy to potential hidden moss diversity.

## Materials and methods

### Accessions and taxonomy

Each specimen is discussed and presented in phylogenetic analysis using its DNA accession number to support future taxonomic work (Supplementary Table S1). This 4- digit identifier is consistent among vouchers, DNA extractions, and sequence data submitted to NCBI’s GenBank. Medina et al. (2019) did not include in their nuclear target capture phylogeny for the 14 samples present in the organellar plastome phylogeny (Medina et al., 2018) though the data were published (https://www.ncbi.nlm.nih.gov/bioproject/PRJNA674709). These 14 samples were initially flagged as allopolyploid due to 1) high numbers of paralog warnings during sequence assembly with HybPiper 1.3.1 (Johnson et al., 2016), and 2) high mean heterozygosity across all loci (>10 bp/locus) in gametophyte tissue estimated by mapping reads to a *Physcomitrium patens* reference sequence. Gametophytes of non-polyploid mosses are haploid and thus are expected to have no or only negligible (i.e. less than 5% of sites, then attributed to sequencing error) detected heterozygosity. Thirteen of the 14 putatively allopolyploid samples are included here in a novel phylogenetic analysis to identify progenitor lineages of these hybrids. One putative allopolyploid is excluded as subsequent analysis revealed poor quality data. In addition, we introduced one accession of *P. collenchymatum,* to address conflicting reports of hybrid status in previous publications. As a result, the final sampling includes 60 accessions comprising 46 haploids from Medina et al. (2019), 13 putative allopolyploids, and one additional sample of *P. collenchymatum.* We identified all samples to their closest morphological species concept using the most appropriate regional floras (e.g., Frey et al., 2006 McIntosh, 2007; Goffinet, 2007).

The newly included sample, labeled *Physcomitrium collenchymatum* (5274A), is collected from a propagated culture (Culture accession number 40061) from the International Moss Stock Culture (IMSC, Freiburg, Germany) that served as the exemplar for this species in prior studies (McDaniel et al., 2010; Beike et al., 2014; Ostendorf et al., 2021). Procedures for targeted sequencing of 806 nuclear loci for this sample followed Medina et al. (2019) with the following differences: we extracted genomic DNA using a modified CTAB/chloroform method, sheared it with the Fragmentase enzyme (New England Biolabs, Ipswich, MA, USA), and prepared shotgun sequencing libraries with the NEBNext Ultra-II DNA kit modified to use half-volumes compared with manufacturer recommendations. We pooled the new sample with other mosses not included in the present study for target enrichment using the MyBaits v3 protocol (Arbor Biosciences, Ann Arbor, MI, USA) and sequenced 2×300 paired-end reads on an Illumina MiSeq at the Texas Tech University Center for Biotechnology and Genomics.

### Sequence Assembly

We recovered targeted sequences from the 13 allopolyploid samples omitted by Medina et al. (2019) in their nuclear phylogenetic analyses, using HybPiper v 1.3.1 (Johnson et al., 2016), as well as from the new sample of *P. collenchymatum.* For each sample, we extracted supercontig sequences, representing exons and flanking non-coding regions for each locus. We called variants within each sample using the GATK (version 4.1.0.6) Variant Detection Workflow (Van der Auwera & O’Connor, 2020), which began by mapping reads from a sample to the recovered supercontigs from the same sample. We used MarkDuplicates (Picard Toolkit, 2019) to identify and remove duplicate reads via common mapping location prior to variant calling with HaplotypeCaller. We saved only the single nucleotide polymorphisms (SNPs) and used a hard filter to retain only SNPs with high mapping quality, read depth, and base call quality (QD < 5.0 && FS > 60.0 && MQ < 40.0 && MQRankSum < -12.5 && ReadPosRankSum < -8.0).

For samples with an F-statistic greater than 0.5, indicating higher than expected heterozygosity, we phased homeologs within each locus using a method similar to that of Kates et al. (2018). We used WhatsHap (version 1.1, Martin et al. 2016) to phase variants and extracted phased sequences at each locus using the Python script haplonerate.py (github.com/mossmatters/phyloscripts). The latter script is necessary to sort sequences within the largest phaseable block (Figure 2), that is the largest stretch of sequence within a locus where variants can be phased using the sequence reads. Our method differs from Kates et al. (2018) which retained variants outside the largest block as ambiguous sites, as we chose the more conservative approach of deleting all sequences that fell outside the largest block. This is consistent with the approach taken by Tiley et al. (2021).

**Figure 2:**
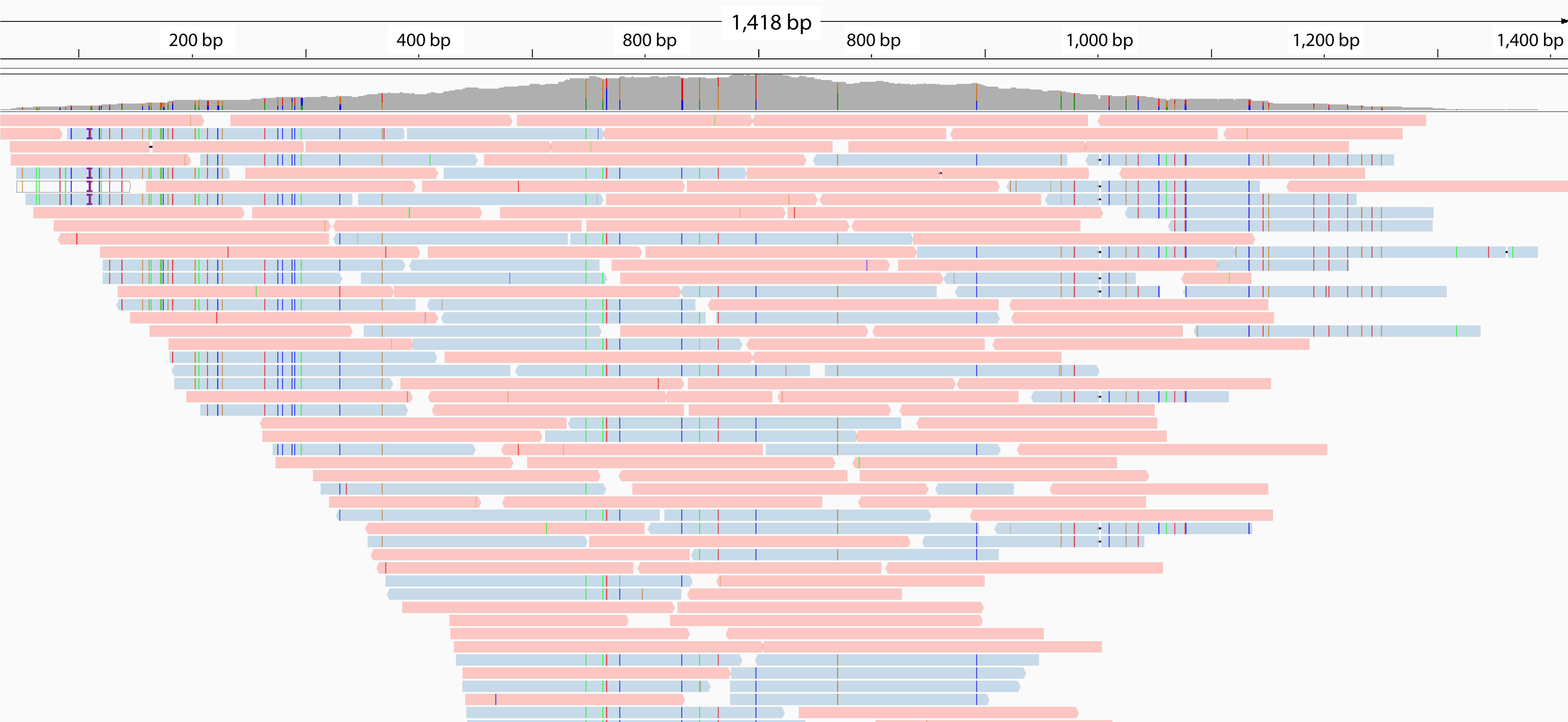
Reads associated with nuclear marker 7379 from *P. immersum-*3176 indicating SNPs in a subset of reads representing two distinct subgenomes in the allodiploids. This is representative of the readbackedphasing approach.

To determine the likely ploidy for our samples with a high F-statistic, we used nQuire (accessed from GitHub 2022-01-01, Weiß et al., 2018) to fit gaussian mixture models to within-sample allele frequencies. In nQuire documentation, sequence variants associated with a locus are referred to as “alleles” though in the present study, when multiple “alleles” are detected, these would be homeologs. This approach estimates the frequency of SNPs with multiple allelic variants. Various allelic ratios (e.g., 0.5/0.5 for diploids) are generated for each ploidy. Mosses are haploid dominant organisms, and hence a non-polyploid individual will have haploid gametophytic tissue and diploid sporophytic tissues (Figure 1). Here, we have sampled and sequenced gametophytic tissue and accordingly expect that polyploids will have a ploidy of two or higher. Because nQuire does not have a model for haploid samples, we were only able to analyze our 14 putatively allopolyploid samples to choose among diploid, triploid, and tetraploid models via maximum likelihood. For all samples, we implemented denoising within nQuire to eliminate low quality reads with noise generated from repetitive sequences.

### Subgenome phasing

We completed the GATK+Whatshap workflow for the 648 genes used in Medina et al. (2019) for the 14 polyploid samples, resulting in reads assembled into homeologs for each locus (Figure 2). Any sequence outside the largest phase block within a locus was deleted from the polyploid samples. As a full Bayesian inference method, computational limitations are a concern, so we chose to phase subgenomes using 50 nuclear loci. We first sampled two subsets of these genes (25 genes each) and analyzed them independently using homologizer. One accession (*P. pyriforme-*3798) has missing data for one of the 50 genes selected, gene number 8583 (Supplementary Table S2). Each locus yielded consistent topologies and placement of each allopolyploid subgenome, suggesting that a subset of single copy nuclear loci likely yields a phylogeny representative of the full dataset. In a homologizer analysis that we will refer to as *iteration one,* we added two phased homeologous sequences per polyploid sample (28 alleles) to the haploid nuclear marker alignments (46 sequences) from Medina et al. (2019) resulting in 74 sequences for each gene. We added homeolog sequences from our 14 putative allopolyploid samples to the trimmed supercontig alignments from Medina et al. (2019) using MAFFT v 7.407 with the addsequence and keeplength options (Katoh& Frith, 2012). Homologizer (Freyman et al., 2023) is a function in the software RevBayes (version 1.1, Höhna et al., 2016) that jointly infers the subgenome phase across loci along with the phylogeny from a matrix of DNA sequences. We followed the methods available from github.com/wf8/homeolog_phasing to prepare a RevBayes control file containing prior associations between homeolog sequences and subgenomes for each sample, proposal frequencies for swapping homeologs between subgenomes, and standard priors for a GTR substitution model. We ran RevBayes for 10000 generations (representing over 5.32 million Metropolis-Hastings moves, Höhna et al., 2016) on four independent runs and checked for chain stationarity in each run via a stable posterior probability and an effective sample size (ESS) of > 200 (a convention for Bayesian inference, Drummond& Bouckaert, 2015) for all parameters using Tracer v1.7 (Rambaut et al., 2018).

In order to attempt to represent polyploids greater than diploid with more than two tips, as we did in *iteration one,* we implemented *iteration two,* or the “fixed-diploid” approach, as follows. For populations with a ploidy greater than two, as estimated by nquire, variants were inferred with GATK changing the --ploidy flag where appropriate. We used the “polyphase” function in whatshap to generate phased haplotypes within each locus. Because earlier runs struggled with convergence when including higher-ploidy samples, we took a tiered approach to phased subgenomes of the higher ploidy samples: first, we used the output of the first iteration to set (fix) the subegenome phasing for diploids across loci in a second run of homologizer, where we only inferred phase for the samples with higher ploidy (*Physcomitrium pyriforme* samples 3410 and 3798). Finally, we implemented a third apporach, *iteration three,* or the “dummy-tip” analysis, for the incorporation of higher ploidies represented by more than two tips. The “dummy tip” method has previously been employed for higher ploidy sequences (Freyman et al., 2023), where a blank sequence was inserted for a high-ploidy sample as a way to accommodate unsampled or lost homeologs. We ran ten independent runs of homologizer for 5000 generations for both the “fixed diploid” and “dummy tip” methods.

For all analyses, in addition to the Bayesian phylogeny inferred using RevBayes, we inferred a maximum quartet species tree using ASTRAL-III (Zhang et al., 2018). Using the joint posterior probability (the posterior probability determined jointly among loci) for subgenome phase from homologizer, we modified the sequence labels for all 14 polyploid samples to indicate assigned subgenomes. We inferred gene trees for each locus, with the subgenomes treated as their own tips, using IQTREE (version 2.0.3, Minh et al., 2020), with substitution models selected using ModelFinderPlus (Kalaanamoorthy et al. 2017) and assessed support based on 1000 UltraFast Bootstrap (Hoang et al., 2018) replicates. We collapsed gene tree branches with less than 30% support using the nw_ed function in newick_utils (accessed from GitHub 2020-06-01, Junier et al., 2010). After inferring a species tree from the collapsed gene trees with ASTRAL-III (version 5.6.3, Zhang et al. 2017), we used phyparts (version 0.0.1, Smith et al. 2015) to assess bipartitions among the 50 genes and visualized concordance and conflict with phypartspiecharts.py and minorityreport.py (github.com/mossmatters/phyloscripts). To determine the extent to which conflict among gene trees could be attributed to the inclusion of allopolyploid subgenome sequences, a second ASTRAL tree including only haploids in the dataset was generated.

### Maternal and paternal subgenomes

Each of the allopolyploid samples in the present study, with the exception of the newly sampled *P. collenchymatum-*5274A, was included in the organellar phylogenomic reconstruction by Medina et al. (2018) and hence should have one nuclear subgenome in a phylogenetic position identical to that in the plastid exome phylogeny reconstructed from the same samples (Medina et al., 2018). These relationships are therefore used here to infer the maternally inherited subgenome in each allopolyploid.

For one sample, *P. pyriforme – 3410*, substantial gene conflict in the positioning of its subgenomes indicated a need to rule out contamination and ensure that *P. pyriforme – 3410* in the present study was the sample for which plastid DNA was extracted and sequenced by Medina et al. (2018). Four plastid genes were recovered from target capture data generated for *P. pyriforme – 3410* generated by Medina et al. (2019) using HybPiper v 1.3.1 (Johnson et al., 2016). Consensus sequences were aligned with single gene alignments from Medina et al. (2018) to assess sequence similarity with *P. pyriforme – 3410*.

### Data availability

All nucleotide alignments, RevBayes control files, gene trees, and species trees are available via the Dryad Digital repository (https://datadryad.org/stash/share/WWVIuGH9Iko4wjVg9SGe2bSNYRWl1WINkuIhbXw 54Hk). For target capture baits used, see Medina et al. (2019) and Dryad Digital repository (https://datadryad.org/stash/dataset/doi:10.5061/dryad.8rq9465).

### Physcomitrium collenchymatum *sequencing*

We tested whether conflicting reports of allopolyploidy in *P. collenchymatum* were attributable to culture contamination, by comparing a barcoding locus sequenced from DNA extracted from the culture propagated in our laboratory (5274A), the original IMSC culture (40061), and the herbarium voucher from which the later was established. In addition, two plastid markers are sequenced for *P. collenchymatum-*5274A and used as the basis of maternal subgenome identity.

#### Sanger Sequencing

For target-capture sequencing of the nuclear loci 4780 and 7379 both homeologs were consistently recovered in allopolyploids. Accordingly, Sanger sequencing was used to generate reads for nuclear markers 4780 and 7379 (Supplementary Table S3) for the IMSC culture (40061) and our associated culture (5274A) of *P. collenchymatum* to confirm their identity (Supplementary Table S1). Nuclear sequences associated with the two cultures were compared with the sequences generated from the original dry voucher specimen (Homberg 1155 MO). In addition, two plastid markers, *psba-*trnH, and *rps*5-trnS, were sequenced for the culture *P. collenchymatum—*5274A to discern the maternal subgenome on the basis of phylogenetic relatedness with haploid samples (Supplementary Figure S2).

#### DNA Extraction and PCR Amplification

For live cultures, total genomic DNA was extracted using the NucleoSpin Plant II Midi kit (Macherey-Nagel. Düren, Germany) following the manufacturer’s instructions. Nuclear markers 7379 and 4780 as well as plastid markers *psb*A-trnH and *rps*5-trnS were amplified using primers detailed in Supplementary Table S3. For the nuclear markers, PCRs were performed in a final volume of 25 µL with 12.5 µL of GoTaq Green Mastermix (Promega, Madison WI, USA), 9.5 µL of water, 1 µL of the forward and reverse primer (10 µm), and 1 µL of genomic DNA extract. PCR consisted of a 3 min hot start at 94°C, followed by 40 cycles of denaturation (1 min, 94°C), annealing (1 min, 50°C) and extension (1 min, 70 °C), ended by a final extension step of 10 min. For plastid markers *psb*A-trnH and *rps*5-trnS, a nested PCR was used. Both PCRs consisted of a 3 min hot start at 94 °C, followed by 40 cycles of denaturation (1 min @ 94 °C), annealing (1 min @ 50°C) and extension (1 min @ 70°C), ended by a final extension step of 10 min. PCR products were cleaned used the ExoSAP-IT protocol (USB-Affymetrix, Cleveland OH, USA) and sequenced at Eurofins Genomics on an ABI prism sequencing platform (Louisville, KY).

## Results

### Within-locus phased assembly

Haploid samples are evidenced by a low proportion of heterozygous sites identified by GATK (Supplementary Table S1) and are represented in phylogenetic analyses with only one sequence per locus. Among the allopolyploid samples previously excluded from the Medina et al. (2019) nuclear phylogeny, the number of variant sites per locus within a sample ranged from 42.86 *(E. hungaricus-*3177) to 133.74 (*P. pyriforme – 3410*) sites per gene, compared to a range of 0.00 to 7.62 for the haploid samples. The samples with high numbers of within-sample variants include four samples of *P. eurystomum,* four of *P. sphaericum,* two of *P. pyriforme,* one of *P. immersum,* two of *Entosthodon hungaricus,* and one of *P. collenchymatum* (Supplementary Table S1).

### nQuire

On the basis of a high F-statistic, diploidy is the best-fit model for 12 hypothesized allopolyploids, whereas triploidy is the best fit model for *P. pyriforme – 3410* and tetraploidy is the best fit for *P. pyriforme*-3798. (Supplementary Table S4; ploidy assessments are based on the gametophyte, hence the nQuire results suggest that *P. pyriforme*-3798 has octoploid sporophytes, see Figure 1). It should be noted that while nQuire provides a hypothesis for ploidy, it is designed for polyploid detection in angiosperms where the dominant generation is diploid, rather than haploid in mosses. Here the dominant haploid generation, the gametophytic phase, is sampled.

### Subgenome phasing and phylogenetic relationships

The apparent triploid and tetraploid, *P. pyriforme – 3410* and 3798, are treated as diploids in *iteration one* of our analyses and as higher-level ploidies (three subgenomes) in *iteration two* (the “fixed diploid” approach) to test homologizer’s capacity to phase higher level ploidies and determine future approaches to improve its robustness. To evaluate the effect of treating *P. pyriforme – 3410* and 3798 as diploids, in iteration one, on inferred phylogenetic relationships among the allodiploid subgenomes, a RevBayes tree excluding *P. pyriforme – 3410* and 3798 is included (Supplementary Figure S1). The topology is largely consistent with that of the ASTRAL tree including all subgenomes (Supplementary Figure S2). The exception is the circumscription of subgenomes associated with *E. hungaricus-*3838 and 3177, since the maternal subgenome of each is sister to the maternal subgenome of triploid *P. pyriforme – 3410* in the iteration one analysis (Supplementary Figure S2). As a result, the RevBayes tree excluding the higher-level ploidies resolves all subgenomes of *E. hungaricus-*3838 and 3177 in one clade on a short branch (Supplementary Figure S1). The remainder of the allopolyploid samples are apparent allodiploids (Supplementary Table S3). The total length of the trimmed alignments for 50 phased nuclear supercontigs was 187988 bp (per gene average = 3759 bp, range 1668 – 12446 bp). The iteration two analysis testing the incorporation of greater than two subgenomes per sample represents *Physcomitrium pyriforme –* 3410, 3798 with three tips, or three subgenomes. The topologies resulting from the ASTRAL tree for the second iteration is largely consistent with the ASTRAL tree from the first iteration representing these populations with two subgenomes. The topologies differ in the position of the likely paternal subgenome of *P. pyriforme –* 3798, and paternal and maternal subgenomes of *P. pyriforme – 3410*. When treated as triploids, *P. pyriforme – 3410* includes two subgenomes in a clade with a subgenome of *P. pyriforme –* 3798 sister to the *P. pyriforme* species complex, and one subgenome sister to the *P. japonicum* and *P. pyriforme* species complex. The remaining two subgenomes of *P. pyriforme –* 3798 are in the *P. pyriforme* species complex (Supplementary Figure S4).

Homologizer yielded two main outputs: 1) a Bayesian phylogenetic inference of phylogeny (including the haploid samples and the subgenomes for each allopolyploid sample) and 2) posterior probabilities for assigning homeologs of each gene to a subgenome. After discarding the first 25% of the run as burn-in, we calculated a maximum credibility tree that shows full support at nearly all nodes for iteration one (Supplementary Figures S5&S6) and iteration two (Supplementary Figure S7). Convergence and stationarity of homologizer runs with all polyploid samples coded as diploid (iteration one) was confirmed using ESS > 200 for all parameters and a stable posterior probability after burnin. By contrast, homologizer runs struggled to reach convergence when the two higher-ploidy samples (*P. pyriforme* 3410 and 3798) were represented by three subgenomes and phased simultaneously with allodiploids (iteration two). For the “fixed diploid” analysis which fixed subgenome phase for diploid samples while we attempted to phase the remaining two samples as triploid, only seven of 10 runs converged within 5000 generations. Finally, for iteration three, the “dummy tip” analysis where a fourth blank sequence was added for the two higher ploidy samples, each of the 10 runs converged on a different posterior probability (Supplementary Figure S8).

Phylogenetic inferences in iterations one and two resolve a tree topology largely consistent with Medina et al. (2019) (Supplementary Figure S6; S7). As before, *P. pyrifome* is resolved as paraphyletic. In iteration one, only two nodes are not resolved with a 100% posterior probability, and the posterior probability of subgenome assignment was maximal (PP = 1.0) for all samples and loci, which allowed assigning each homeolog to a subgenome for further analysis (Supplementary Figure S6).

The subgenomes of *Entosthodon hungaricus-*3838 and 3177 are the two most phylogenetically distant subgenomes in the sampling: the maternal subgenome belongs to a lineage sister to *Physcomitrium* and the paternal subgenome originated from the *E. attenuatus* clade (Figure 3). The paternal subgenomes of *E. hungaricus-*3828 and 3177 are resolved as part of the *Entosthodon* clade with minimal genic conflict (39/50 genes are concordant). However, gene conflict (only 20/50 genes are concordant) is substantial at the ancestral node of the clade including the maternal subgenomes of *E. hungaricus-*3828 and 3177. As in the ATSRAL phylogeny generated in iteration one, in iteration two, there is little genic conflict in the resolution of the paternal genomes, and substantially more conflict in the resolution of the maternal genomes (Supplementary Figure S7).

**Figure 3.**
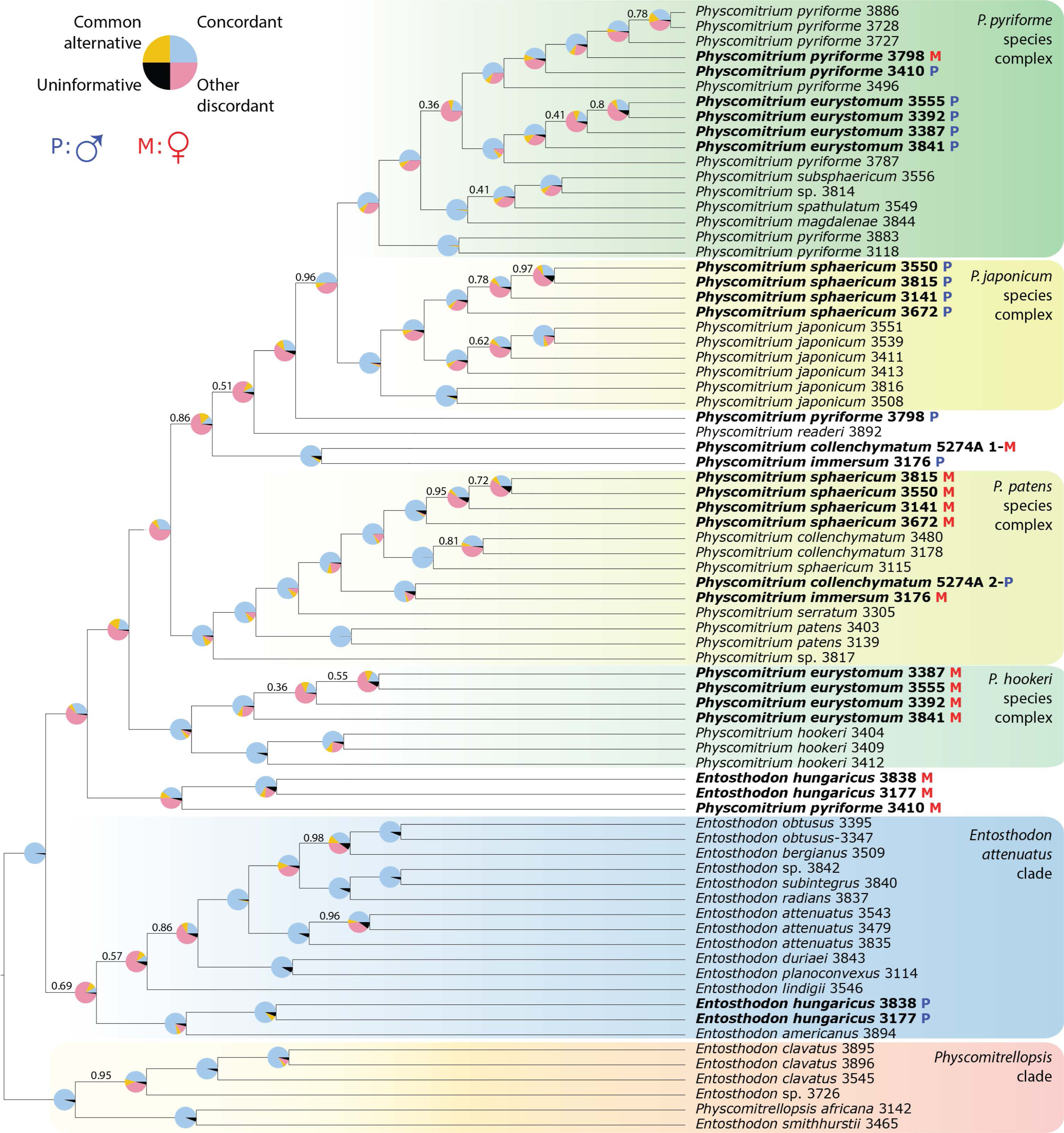
Phylogenetic reconstruction of haploid samples and phased subgenomes of allodiploids using ASTRAL from 50 gene trees. Pie charts on each branch indicate the proportion of gene trees concordant with that bipartition (blue), proportion of gene trees with the most common discordant bipartition for that bipartition (orange), proportion of gene trees with all other discordant bipartitions for that bipartition (red) and proportion of gene trees with no support due to missing data for that bipartition (black). Support values are local posterior probability from ASTRAL; only values less than 1.0 are shown. Boxes indicate named clades as discussed in text. Taxa in bold are allopolyploids represented by their maternal (M) and paternal (P) subgenomes.

The specimens *Physcomitrium sphaericum-*3141, 3672, 3550 and 3815 are from China and their paternal subgenome is from the *P. japonicum* species complex, whereas the maternal subgenome is from the *P. patens* species complex related to (haploid) *P. collenchymatum.* The respective subgenomes of *P. immersum-*3176 and *P. collenchymatum-*5274A are sister to each other, with one parental subgenome sister to the clade comprising *P. readeri*, the progenitor to the *P. japonicum* and *P. pyriforme* species complexes and the second sister to a clade including *P. collenchymatum* and *P. serratum.* (Figure 3; Supplementary Figure S7). However, the maternal and paternal contributions of the two proposed progenitors to these two allopolyploid lineages are reversed with the maternal subgenome of *P. immersum-*3176 in the *P. patens* species complex and the maternal subgenome of *P. collenchymatum-*5274A sister to the *P. pyriforme* species complex (Figure 3). *Physcomitrium eurystomum*-3841, 3392, 3387 and 3841 comprise two subgenomes, with the paternal parent closely related to *P. pyriforme*-3787 and the maternal one sister to *P. hookeri* (Figure 3; Supplementary Figure S7).

In the iteration one analysis our ASTRAL analysis of gene trees inferred from phased subgenome sequences revealed a topology largely congruent with the concatenated topology from RevBayes (Figure 3; Supplementary Figures S5&S6). The only partially supported incongruence pertains to the placement of the paternal subgenome of *P. pyriforme*-3798: in the RevBayes tree it is nested in a clade with the maternal genome of *P. pyriforme – 3410*, whereas in the ASTRAL tree its placement is unsupported along the backbone of *Physcomitrium*. Other disagreements, including the placement of *P. readeri*, are unsupported in the ASTRAL analysis. Short internodes in part indicate a high level of conflict among the 50 genes, particularly for the monophyly of the clade comprising the *P. pyriforme* and *P. japonicum* species complexes and for the placement of *P. readeri* and subgenomes of *P. immersum* and of the polyploid *P. collenchymatum* (Figure 3).

Bipartition analysis using phyparts (Smith et al., 2015) reveals some incongruence among genes supporting relationships of allopolyploid subgenomes to haploid samples. For example, in the iteration one analysis, which treats *P. pyriforme – 3410*, as a diploid, the species tree placement of the maternal subgenome is resolved with the *E. hungaricus* maternal subgenome by 20 of 50 genes. The placement of the paternal subgenome of *P. pyriforme – 3410* with haploid samples of *P. pyriforme* from North Carolina in the *P. pyriforme* species complex is supported by 22 genes. In both cases, conflict arises from substantial minority bipartitions: nine genes reconstruct the two homeologs from sample *P. pyriforme – 3410* as sister to each other, while the homeologs from the two *E. hungaricus* samples form a clade in 15 gene trees (Supplementary Figure 2). Similarly, in the iteration two analysis, i.e. “fixed diploid” approach including three subgenome tips for *P. pyriforme – 3410*, 3798, the nodes supporting placement of each subgenome is supported by at most 15 genes (Supplementary Figure S4).

By contrast, the placement of *P. immersum* subgenomes sister to those of the allodiploid sample of *P. collenchymatum* (5274A) is consistent for the paternal (iteration one: 45 gene trees; iteration two: 47 gene trees) and maternal subgenomes (iteration one: 39 gene trees; iteration two: 4 gene trees) with no dominant minority bipartition (Supplementary Figure S2; Supplementary Figure S4). For the *P. sphaericum* samples from China, homeologs from the maternal subgenome consistently form a clade (iteration one: 41 gene trees; iteration two: 47 gene trees) sister to (haploid) *P. collenchymatum*. While more incongruences exist among genes for the placement of paternal homeologs of *P. sphaericum* (41 gene trees), their placement within a clade of *P. japonicum* has high support (30 gene trees). Similarly, the relationships of *P. eurystomum* paternal and maternal subgenomes are supported by 42 congruent genes, respectively (Supplementary Figure S2; Supplementary Figure S4).

### Sanger sequencing results

The original culture of *P. collenchymatum* from IMSC (40061), our own culture (5274A), and the original voucher specimen (Homberg 1155, MO) associated with these cultures, share identical sequences for two nuclear markers, each with the same signatures of heterozygosity in each sample (Supplementary Figure S3). In addition, phylogenetic affinities of the plastid sequences *psb*A-trnH and *rps*5-trnS generated from both cultures of *P. collenchymatum* indicate the maternal subgenome is sister to the *P. pyriforme* species complex (Supplementary Figure S9). Assembled chromatograms are available on Dryad (https://datadryad.org/stash/share/WWVIuGH9Iko4wjVg9SGe2bSNYRWl1WINkuIhbXw 54Hk).

### P. pyriforme – 3410 identity

Four plastid genes were recovered from target capture data generated for *P. pyriforme – 3410* by Medina et al. (2019) using HybPiper v 1.3.1 (Johnson et al., 2016). Consensus sequences were aligned with single gene alignments from Medina et al. (2018) to assess similarity with *P. pyriforme – 3410* via target capture of organellar exons for that study. The sequences recovered from target capture data associated with *P. pyriforme – 3410* used in the present study were identical to plastid sequences published for *P. pyriforme – 3410* in Medina et. al. (2018), confirming that the same sample is analyzed in the present study (Supplementary File S1).

## Discussion

We find evidence of allopolyploidy across multiple populations of several species in the *Entosthodon-Physcomitrium* complex, suggesting independent hybridization events in the evolutionary history of the group. We confirm the hybrid origins of *P. eurystomum, P. collenchymatum,* and *E. hungaricus* (McDaniel et al., 2010; Beike et al., 2014; Ostendorf et al., 2021) and reveal the allopolyploid nature of other accessions corresponding to established phenotypically distinct species *P. immersum, P. pyriforme,* and *P. sphaericum.* This analysis allowed us to explore the limitations of the subgenome phasing approach, homologizer, relative to other similar methods. For instance, *P. pyriforme – 3410*, 3798 are the first proposed triploid and tetraploid accessions identified in the *P. pyriforme* species complex using a molecular approach, but the accurate circumscription of their subgenomes remains a challenge, particularly given the need to accurately phase subgenomes. The present study contributes to a growing body of evidence of polyploidy and hybridization contributing to the evolution and diversification of the Funariaceae and provides evidence for overlooked diversity. Furthermore, our approaches to homeolog and subgenome phasing are novel in that homologizer is the first tree-based method that phased gene copies simultaneously using all loci simultaneously. The application of a subgenome phasing approach using target capture data is novel for mosses, which is of particular importance since the haploid phase is sampled. As a result, this study lays the foundation for rapidly developing computational approaches to elucidating reticulate species complexes rather than excluding polyploid taxa when inferring phylogenies.

### *Evolutionary origins of known allopolyploids in* Physcomitrium *and* Entosthodon

The evidence of allopolyploid origin of *E. hungaricus, P. eurystomum,* and *P. collenchymatum* according to previous hypotheses (McDaniel et al., 2010; Beike et al., 2014; Ostendorf et al., 2021) is here significantly strengthened when extended to 50 loci. Further, the ability to phase the subgenomes and estimate their phylogenetic affinities within a broader taxonomic sampling (Medina et al., 2018; 2019) revealed novel hypotheses regarding their progenitor species.

Difficulties in discerning the evolutionary origins of the European endemic *E. hungaricus* suggest that further taxon sampling may be needed to identify progenitors. The two *E. hungaricus* populations sampled here are from Austria and Hungary. The paternal subgenome of both samples is sister to the North American *E. americanus* in the *E. attenuatus* clade, whereas its maternal subgenome is sister to a clade of North American lineages (Figures 3 & 4; Supplementary Figures S4 & S7). The phylogenetic placement of each paternal subgenome is well resolved with little conflict among component gene tree topologies (Figure 3; Supplementary Figure S4). Given that the maternal subgenome of *E. hungaricus* is sister to a subgenome of an allopolyploid population of *P. pyriforme* in iteration one (Figure 3), the two allopolyploids may share one progenitor not sampled here or that is extinct (Karbstein et al., 2022). It should be noted that the clade including the maternal subgenomes of allopolyploid *E. hungaricus* is resolved sister to the *Physcomitrium* clade with substantial genic conflict in iteration one (Supplementary Figure S2) and ASTRAL phylogenetic inference excluding *P. pyriforme – 3410* subgenomes results in the resolution of the maternal subgenomes in the *Entosthodon* clade (Supplementary Figure S1). This suggests that the *P. pyriforme – 3410* subgenomes were incorrectly phased. In iteration two of homologizer representing *P. pyriforme – 3410* with three subgenomes, none of the tips remain sister to a subgenome of *E. hungaricus –* 3838, 3177.

**Figure 4.**
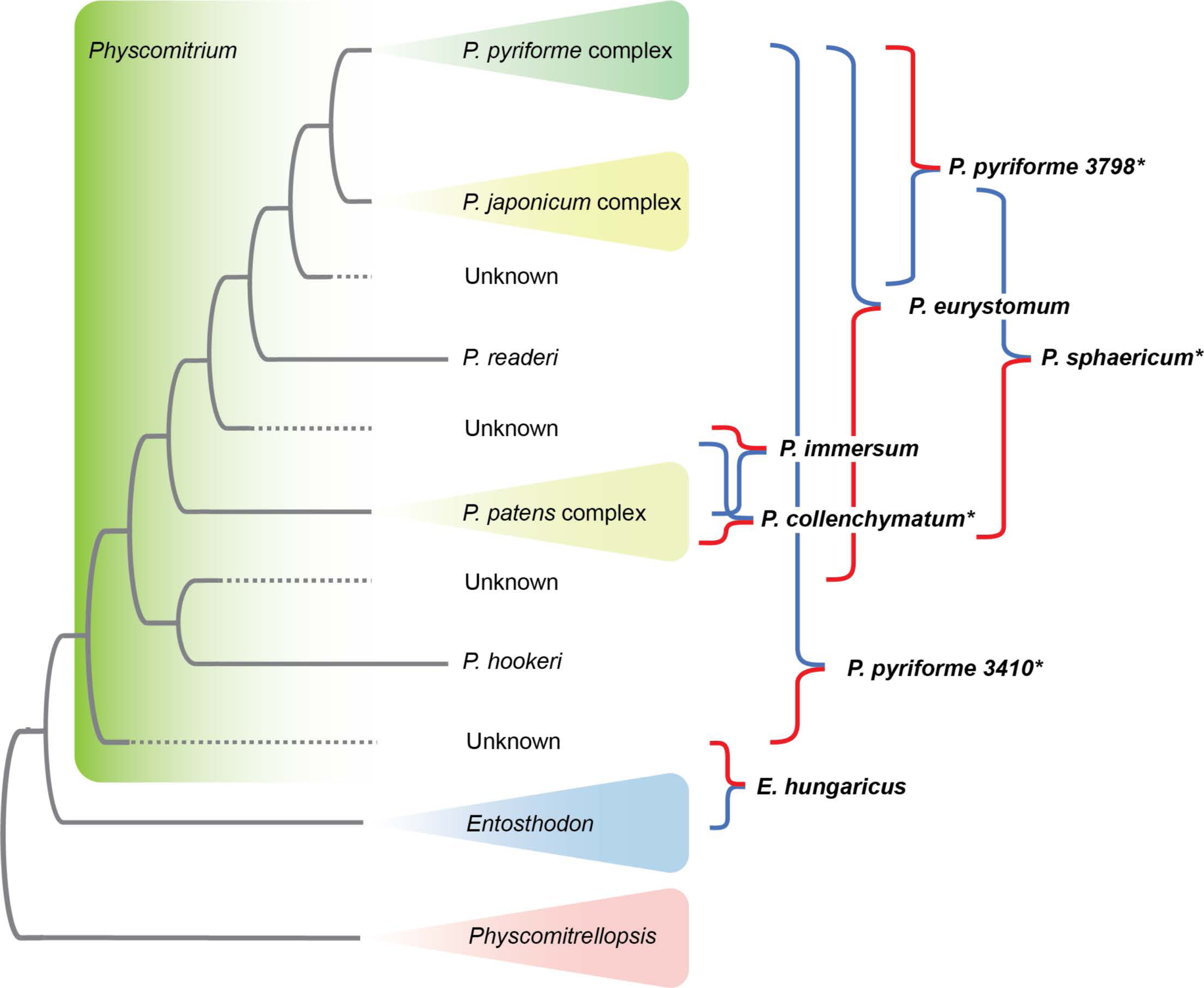
Phylogenetic relationships among major clades represented in Figure 3, inferred using ASTRAL based on 50 genes. Allopolyploid accessions are indicated with brackets adjacent to the phylogeny. Red and blue lines indicate maternal and paternal heritage, respectively. Taxon names currently applied to allopolyploid and haploid accessions are marked with an asterisk.

The two subgenomes of *P. eurystomum* are resolved sister to *P. hookeri* and one population (3787) of the polyphyletic *P. pyriforme,* respectively. Neither parental subgenome is closely related to *P. sphaericum* as previously suggested (McDaniel et al., 2010; Beike et al., 2014; Ostendorf et al., 2021) although our sampling does not include European populations of the latter species. This morphospecies includes haploid and hybrid populations and neither subgenome of the latter forms a clade with the haploid sample (Figure 3; Supplementary Figure S4; see below).

Populations morphologically matching the North American endemic *P. collenchymatum* are either haploid (Medina et al., 2018; 2019) or allopolyploid (McDaniel et al., 2010; Beike et al., 2014; Ostendorf et al., 2021). Sampling numerous nuclear loci for the IMSC stock culture (accession number 40061) of *P. collenchymatum-*5274A, which was used in the latter studies, confirms extensive heterozygosity and hence allopolyploidy (Supplementary Table S1). Furthermore, heterozygosity in the nuclear barcoding loci 4780 and 7379 (Supplementary Figure S3) confirms that the original collection serving as voucher of the IMSC culture (i.e., Homberg 1155), and which is morphologically consistent with *P. collenchymatum,* is an allopolyploid. Consequently, the *P. collenchymatum* morphotype is represented by two distinct lineages, one haploid and one allopolyploid, the latter resulting from a hybridization event involving a haploid species of the *P. patens* species complex and a yet unidentified haploid lineage sister to the clade comprising *P. readeri*, the progenitor to the *P. japonicum* and *P. pyriforme* species complexes (Figure 3). These parental lineages also gave rise via hybridization to *P. immersum* (see below), although with reversed progenitor roles. Whether the name *P. collenchymatum* should apply to the haploid or the allopolyploid lineages must await the characterization on the type specimen of *P. collenchymatum* collected in Missouri (Gier, 1955), as both ploidies are herein reported from this state.

### Newly identified allopolyploid lineage

*Physcomitrium immersum* is here newly identified as another allodiploid species of *Physcomitrium.* Its two subgenomes are highly similar to those of the hybrid accession of *P. collenchymatum-*5274A. However, phylogenetic inferences from plastid data reveal opposite progenitor roles, with one parental lineage in the *P. patens* complex serving as the maternal parent of *P. immersum* and the paternal parent of *P. collenchymatum* (Figure 3; Supplementary Figure S9). Distinct hybrid origins of these species, despite the sister relationships of their respective subgenomes, is consistent with their conspicuously distinct sporophytes (e.g. capsule immersed in *P. immersum* and exserted in *P. collenchymatum*). Repeated hybridization resulting in multiple morphologically distinct allopolyploid lineages is known in ferns and angiosperms (Soltis et al., 2014; Patel et al., 2018; Sessa et al., 2018) and may often represent independent hybridization with inverted parental roles (Sessa et al., 2018).

*Physcomitrium sphaericum* is currently regarded as a haploid species, based on a genotyped population from Germany (McDaniel et al., 2014). In Asia, populations matching this species’ morphotype are either haploid (3115) or allopolyploid (3141, 3550, 3672 and 3815; Figure 3). For the latter, the paternal subgenome is in the *P. japonicum* species complex, whereas the maternal parent may be closely related to the haploid *P. collenchymatum* in the *P. patens* species complex (Figures 3 & 4; Supplementary Figure S4). The haploid and allopolyploid Asian populations of *P. sphaericum* are morphologically indistinguishable. In addition, given strong morphological similarity between *P. sphaericum* and *P. japonicum* (Xing-jiang et al., 2003), if indeed the allopolyploid *P. sphaericum* results from hybridization between these two haploid lineages, one would expect it to be perhaps morphologically indistinguishable from either progenitor.

*Physcomitrium pyriforme* has a long-documented history of polyploidy, with Patel (2021) noting karyotypic evidence for at least five cytotypes (n = 9, 18, 27, 36, 45, 54; see Fritsch, 1991). In the present study, three of the nine *P. pyriforme* populations sampled are allopolyploid, and *P. pyriforme – 3410* and 3798 appear to be triploid and tetraploid, respectively. (Supplementary Table S4). However, we find substantial challenges to inferring progenitors for these higher level (above diploid) polyploids. Although the phylogenetic position of each subgenome of *P. pyriforme* – 3410, 3798 varies depending on the number of subgenomes phased, and with substantial gene conflict, we can reliably infer the identity and correct position of the maternal subgenome using the organellar phylogeny presented in Medina et al. (2018). Plastid phylogenetic inference from Medina et al. (2018) resolves *P. pyriforme* – 3410 as sister to E. hungaricus – 3177, 3838, which is in conflict with all 3410 subgenome positions when represented by three tips in the iteration two (“fixed diploid”) analysis (Supplementary Figure S4), but consistent with the phylogenetic position of the inferred maternal subgenome of 3410 when it is represented by two subgenomes in the iteration one analysis (Figure 3). Still, both analyses indicate substantial gene conflict at nodes ancestral to each 3410 subgenome, suggesting that these conflicts are potentially a result of incomplete lineage sorting or, more problematically, recombination among subgenomes (Oxelman et al., 2017).

While some studies of reticulate evolution phase subgenomes from individuals with ploidies greater than two and incorporate associated subgenomes into phylogenetic analysis, they largely benefit from the use of long-read sequencing approaches such as PacBio, which obviates the necessity of phased sequence assembly (Rothfels et al., 2017; Dauphin et al., 2018). With the short-reads based target capture sequencing employed here and increasingly in phylogenetics (Hale et al., 2020), misassembly and uneven sequencing depth across genes must be contended with and present a challenge to allele phasing in GATK and hence phylogenetic resolution (Eaton et al., 2017; Karbstein et al., 2020). Often, short reads can lead to multiple phased blocks, complicating our ability to phase variants in relationship to variants in other discontinuous blocks (Kates et al., 2018; Slenker et al., 2021). Recent studies leveraging short-read target-capture data in subgenome phasing and parsing allopolyploid origins among plants manage these complications with various approaches including only retaining the largest phase block (Tiley et al., 2021; Slenker et al., 2021). We’ve dealt with the matter of linking variants by using reads-backed phasing and retention of only the largest phased block (Figure 2). Nonetheless, without sufficient read depth, accurate phasing of each subgenome becomes increasingly difficult as ploidy increases (Slenker et al., 2021). Consequently, incorporation of more than two genomes for the triploid and tetraploid samples in our phylogenetic analysis (Supplementary Figure S4), may contribute to the substantial gene conflict at nodes representing divergence of the subgenomes of *P. pyriforme*-3798 and *P. pyriforme – 3410*, respectively. For instance, in iteration two only 11 genes support the relationship of one subgenome of *P. pyriforme – 3410* to the *P. pyriforme* clade (Supplementary Figure S4). Subgenome phasing could be improved with additional target capture data to improve read-depth. Phylogenetic resolution of subgenomes for ploidies higher than two is a critical area of improvement for both read-backed phasing within loci and homologizer and similar target capture-based approaches to subgenomic assignment and phylogenetic analysis.

### Implications of allopolyploidy for moss speciation and diversification

With the inclusion of allopolyploid lineages in taxonomically broad, multi-locus analyses, it is possible to address several core questions in moss evolution. Here, we explore the effect of polyploidization on diversification and trait evolution among polyploid complexes, as well as the extent of hybridization among long diverged lineages.

In some cases, angiosperms polyploidy is associated with a higher rate of speciation (Jiao et al., 2014; Van de Peer et al., 2017; Landis et al., 2018). Others advocate the dead-end hypothesis (Mayrose et al., 2011; 2015). Within the Funariaceae, no two morphologically distinct allopolyploid species share a unique pair of common ancestors with the same progenitor roles; Thus, whereas allopolyploidization may be a frequent speciation mechanism in the Funariaceae, even when involving identical progenitor lineages as in the case of *P. immersum* and *P. collenchymatum,* genome merger does not appear to be followed by subsequent speciation events. However, the lack of diversification may be attributable, in part to the recency of allopolyploid divergence or a need for more dense taxon sampling (Escudero et al., 2014; Mayrose et al., 2014; Dauphin et al., 2018). Relying on maternal node age estimates in Medina et al. (2018) from plastid phylogenetic analysis of populations sampled in the present study, allopolyploid lineages likely arose within the last 5 my (*E. hungaricus*) or even within only the last 1 my (e.g., *P. sphaericum, P. immersum, P. eurystomum*). Given the relatively recent divergence of allopolyploids in the Funariaceae, it is likely that all progenitors are extant. Further, the taxon sampling here is narrow with respect to the 300 species described in the Funariaceae and the population sampling is limited to one or two populations per species. Progenitor hypotheses, including for *E. hungaricus,* would be refined by an expanded taxon and population sampling.

Allopolyploid genomes, by combining homeologs from divergent lineages, are historically thought to present physiological challenges to survival and reproduction (Comai, 2005; Buggs et al. 2008). Accordingly, it is expected that hybridization is less common between more divergent lineages because of the accumulation of phenotypic divergence and hence potential reproductive barriers (Presgraves, 2002; Coyne& Orr, 2004; Moyle et al., 2004). Nonetheless, intergeneric hybrids among lineages that diverged up to 50 mya are known in ferns and angiosperms (Forster & Dale, 1983; Rothfels et al., 2015; Lehtonen, 2018). In our inferences, the parental genomes of each allopolyploid vary in their date of divergence. The most extreme is *Entosthodon hungaricus,* a fertile lineage which combines genomes of parental lineages that diverged approximately 30 mya (Medina et al., 2018). The proposed progenitor genomes of *P. eurystomum, P. sphaericum,* and *P. immersum* diverged less than 15 mya, though here we have not estimated the date of the hybridization event itself.

Since we identified samples based on morphological species concepts, our inferences reveal that at least three morphospecies (i.e., *P. collenchymatum*, *P. pyriforme*, and *P. sphaericum*) comprise both haploid and allopolyploid genotypes, challenging diversity estimates, taxonomic delimitation and nomenclatural assignment. The extent to which morphological features can be used to reliably identify species in the *Physcomitrium-Entosthodon* complex requires further study. The size of spores is correlated with genome size in both ferns and mosses and hence may be valuable in identifying polyploids in a narrow taxonomic context (Košnar et al., 2012; Barrington et al., 2020). Taxonomic treatment in the complex may need to follow the example of *Sphagnum*, in which hybridization has been frequently demonstrated (Ricca & Shaw 2010; Ricca et al., 2011; Karlin & Smouse, 2017). For example, the *Sphagnum subsecundum* complex includes haploid and allodiploid species as well as morphospecies that includes multiple cytotypes. (Shaw et al., 2012). In the *Physcomitrium-Entosthodon* complex, detailed surveys of morphological characters are needed to make similar progress. Because the type specimens of the taxa studied here are 60–120 years old and sometimes consist of a few tiny plants for destructive sampling, estimating ploidy, even based on heterozygosity of barcodes, would be difficult or impossible. Morphological characters may diagnose lineages of distinct ploidy and hence could allow assigning a morphotype to a type specimen and applying the correct name. However, critical morphometric analyses are lacking, and as currently circumscribed morphospecies comprise populations of haploid and hybrid populations.

### Challenges to allopolyploid discovery, homeolog phasing, and homeolog sorting

The inclusion of allopolyploid lineages along with haploid taxa in a phylogeny required two phasing steps: phasing of reads within loci and of homeologs within samples. We initially identified potential allopolyploids via an abundance of “paralog warnings” in HybPiper. We then characterized the degree of heterozygosity after mapping reads to the HybPiper output and identifying within-sample variants with GATK. Our results suggest the paralog warning method was not perfect; several samples, including *E. hungaricus*-3177, did not have many paralog warnings but were nevertheless highly heterozygous when put through the GATK workflow. It is likely that low sequencing depth can confound the ability of HybPiper version 1.3 to identify multiple copies, because it only does so when multiple full-length contigs are recovered for a gene. (Notably, version 2 of HybPiper purports to alleviate this shortcoming and will produce warnings if any part of the assembled sequence has multiple representative contigs; see github.com/HybPiper/wiki). An alternative approach to allopolyploid discovery with target capture data may be to use HybPhaser (Nauheimer et al., 2021) for large-scale review of target-capture data to identify homeologs and to discover hidden paralogs and contaminants.

Our read-backed phasing workflow using GATK and WhatsHap often resulted in fully phased sequences across thousands of bases, suggesting that our nuclear target capture genes have many short introns that our relatively long (300 bp) paired-end Illumina reads could sequence through. The within-locus phasing (assembly) workflow takes advantage of programs written for phasing heterozygosity in diploid-dominant organisms, and whether this will scale easily to higher ploidy remains to be tested. In our multi-tip analysis, we’ve modified our read-backed phasing workflow to incorporate the “polyphase” function in WhatsHap allowing for the phasing of haplotypes at ploidies greater than two (Schrinner et al., 2020). Whereas three distinct haplotypes were phased for *Physcomitrium pyriforme –* 3798 and 3410, substantial gene conflict is evident in the phylogenetic position of each subgenome (Supplementary Figure S8). Detecting autopolyploids and unbalanced allopolyploids such as triploids using this approach may be much more difficult by requiring divergence between subgenomes substantial enough for detection as unique homeologs among multiple genes as well as sufficient read depth across variants (Karlin& Smouse, 2017). This challenge was reflected in the inconsistency in posterior probability across homologizer runs when our two higher ploidy samples were coded as triploids. It is possible that recombination among subgenomes – as has been previously reported in triploid *Sphagnum* mosses (Karlin& Smouse, 2017) - may have occurred since the genome doubling events in the polyploid *P. pyriforme* lineages. This could contribute to lack of convergence in homologizer and the high levels of subgenome discordance observed in the species tree reconstructed from the iteration two “fixed 2N” homologizer results (Supplementary Figure S8).

Newly added features in programs including Whatshap “polyphase” (Schrinner et al., 2020) and H-POP-G (Xie et al., 2016) are promising, but more studies using these features to analyze polyploid target capture data are needed to explore the utility of these tools in confidently calling variants (homeologs in the present study) with lower sequencing depth. Our target capture data were generated from enrichment of genomic DNA extracted from gametophytic tissue exclusively. This simplifies analysis by necessitating only the assignment of alleles to monoploid genomes. Analyzing data derived from sporophyte allodiploids, as we did for those of gametophytes, would be more complicated as it requires allele phasing and subgenome assignment for up to six variants at each locus. This is a key issue in that one major benefit of target capture sequencing is that it can be an effective method for generating subgenomic data even for fragmented DNA derived from herbarium specimens (Viruel et al., 2016; 2019).

Our results rely on subgenome phasing in the RevBayes plugin homologizer to phase among-loci, and we highlight its utility for integration of polyploids in phylogenetic systematic studies. However, the scaling potential of homologizer is limited without future improvements. For instance, with 50 genes (representing about 188 Kb) and 74 tips, RevBayes took over six days to complete 10000 generations on a high-performance computer.

In this study, we observe much more discordance among gene trees as compared to that in Medina et al. (2019). Although phylogenetic inference in Medina et al. (2019) is based on 648 genes compared to 50 in the present study, much of the discordance we observed is likely attributable to the inclusion of phased allopolyploid subgenomes in gene tree construction. This is evident when comparing the phylogeny excluding allopolyploids (Supplementary Figure S10) with the phylogeny including all samples (Figure 3; Supplementary Figure S4), both based on the same 50 genes. Many of the discordant gene trees result from phased subgenome sequences resolved as sister to each other, which suggests that some genes may have more complicated inheritance, i.e., the loss or duplication of one homeolog after polyploidization. Investigating the mode of polyploid inheritance and homeologous exchange would require phasing many more genes than can currently be practically supported by homologizer. One approach could be to conduct iterative phasing by phasing multiple sets of genes in parallel (e.g., 50 sets of 50 genes each), and then using a final phasing of one gene per set to connect all homeologs into subgenomes for final analysis. We also limited our search to genes with complete data for all samples and increasing gene sampling will need to accommodate missing data.

Finally, although target capture sequencing of low and single copy nuclear markers is critical to the identification and phylogenetic analysis of allopolyploids and allopolyploid subgenomes, our study also highlights the importance of highly supported organellar phylogenies to the identification of allopolyploid progenitors. For each polyploid hybrid species, we resolved one subgenome in a position consistent with that of the organellar genome proposed by Medina et al. (2018), except for *P. collenchymatum*-5274A which was not sampled in the previous study. Although the plastid exome likely represents just one coalescent history, it is valuable for identifying the maternal lineage in hybrids and determining whether polyploids have recurrent formation.

## Conclusions

The diversification of the Funariaceae is characterized by significant reticulation among multiple species complexes, and even between genera. The Funariaceae exemplify growing evidence for the significance of polyploidy in moss speciation and evolution (Sawangproh and Cronberg, 2021). Numerous studies (Soltis et al., 2009; Barker et al., 2016; Schmickl et al., 2017) in plants suggest that polyploids constitute a substantial proportion of total plant species diversity (Wood et al., 2009) and hence accurate circumscription of polyploid subgenomes facilitates the testing of questions in evolutionary biology pertaining to diversification. Here, we demonstrate the importance of phasing and sorting allopolyploid subgenomes for phylogenomic analysis. The implementation of homologizer as a tool for the incorporation of allopolyploids in phylogenetic analysis is a critical step toward elucidating the role of allopolyploidy in the evolution and diversification of plants.

## Supplementary

Table S1. For each sample, collector and locality data, the total number of heterozygous sites, number of genes recovered, paralogy for all specimens.

l*psb*A-trnH and *rps5-*trnS.

Table S2. A list of gene names for the 50 nuclear loci analyzed using Homologizer, which were selected from the 648 gene set published in Medina et al. (2019).

Table S3. Primer sequences and citations for all sanger sequences markers for sample identity validation.

Table S4. Results from nQuire analysis of all samples. Values in columns 2-5 are likelihood values indicating fit of a free model and the lrdmodel for diploids, triploids, and tetraploids. Columns 6-8 indicate the delta-likelihood, or change in likelihood, between a free model and the lrdmodel for diploids, triploids, and tetraploids. Values in bold are the lowest likelihoods for each sample.

Figure S1. Phylogeny of the *Entosthodon-Physcomitrium* complex inferred from 50 genes and flanking regions. RevBayes tree including phased allopolyploid subgenomes sampled in the present study, with posterior probability indicated.

Figure S2. Phylogeny of the *Entosthodon*-*Physcomitrium* complex inferred from 648 nuclear genes and flanking regions. ASTRAL tree including all haploids and allopolyploids sampled, with all gene congruence indicated at each node. Pie charts on each branch indicate the proportion of gene trees concordant with that bipartition (blue), proportion of gene trees with the most common discordant bipartition for that bipartition (orange), proportion of gene trees with all other discordant bipartitions for that bipartition (red) and proportion of gene trees with no support due to missing data for that bipartition (black).

Figure S3: Alignment of chromatograms of single copy nuclear marker 7379 for *P. collenchymatum-*5274A and the associated voucher (*Homberg 1155*) and stock culture (IMSC 40061).

Figure S4. Phylogenetic reconstruction of haploid samples, phased diploid subgenomes, and three phased subgenomes of *P. pyriforme – 3410*, 3798 using ASTRAL from 50 gene trees, with allodiploids sungenome phases fixed (“iteration two“ analysis). Pie charts on each branch indicate the proportion of gene trees concordant with that bipartition (blue), proportion of gene trees with the most common discordant bipartition for that bipartition (orange), proportion of gene trees with all other discordant bipartitions for that bipartition (red) and proportion of gene trees with no support due to missing data for that bipartition (black). Boxes indicate named clades as discussed in text. Taxa in bold are allopolyploids represented by their maternal (M) and paternal (P) subgenomes.

Figure S5. Phylogeny of the *Entosthodon*-*Physcomitrium* complex inferred from 648 nuclear genes plus flanking regions. ASTRAL tree with branch lengths proportional to concordance.

Figure S6. Phylogeny of the *Entosthodon*-*Physcomitrium* complex inferred from 648 nuclear genes plus flanking regions. RevBayes tree with posterior probability support values. Allopolyploid subgenomes are highlighted with colors unique to individual allopolyploids.

Figure S7. Phylogeny of the *Entosthodon*-*Physcomitrium* complex inferred from 50 nuclear genes plus flanking regions with *P. pyriforme – 3410*, 3798 represented by three subgenome tips (“iteration two” analysis). ASTRAL tree with local posterior probability support values.

Figure S8. Posterior probability trace for Iteration three of homologizer.In this iteration, the phase for allodiploid samples were fixed to the subgenomes from iteration one. The putative allotriploids *P. pyriforme – 3410* and 3798 had three sequences per gene, plus a blank “dummy” sequence to allow for alternative locations of cryptic subgenomes. The trace summarizes posterior probability across 10 independent runs of RevBayes- homologizer, with 5000 generations each. There is a general lack of convergence both within runs and across runs.

Figure S9. Bayesian inference phylogeny based on plastid markers psbA-trnH and *rps*5- trnS. Sampling includes a subset of accessions included in Medina et al. (2019) and *P. collenchymatum*-5274 from the present study and is used to infer the maternal heritage of allopolyploid accessions of *P. collenchymatum* presented in Figure 3.

Figure S10. Phylogeny of the *Entosthodon*-*Physcomitrium* complex inferred from 648 nuclear genes and flanking regions. Astral tree including only haploids sampled in the present study, with all gene congruence indicated at each node. Pie charts on each branch indicate the proportion of gene trees concordant with that bipartition (blue), proportion of gene trees with the most common discordant bipartition for that bipartition (orange), proportion of gene trees with all other discordant bipartitions for that bipartition (red) and proportion of gene trees with no support due to missing data for that bipartition (black).

File S1. Sequences of plastid genes extracted from target capture data for *P. pyriforme – 3410*. They are aligned with plastid sequences from Medina et al. (2018) associated with the same population in order to confirm the identity of this population in the present study.

## Acknowledgements

We thank the IMSC for providing cultures of *Physcomitrium collenchymatum* and John Atwood and Bruce Allen (MO) for the loan of the original vouchers of these cultures.

This research was supported by the National Science Foundation with grants to BG (NSF-DEB 1753811), RM (NSF-DEB 1753673), and MGJ (NSF-DEB 1753800). We thank C. Rothfels and W. Freyman for assistance running Homologizer.

## Data Accessibility

All nucleotide alignments, RevBayes control files, gene trees, and species trees are available via the Dryad Digital repository (https://datadryad.org/stash/share/WWVIuGH9Iko4wjVg9SGe2bSNYRWl1WINkuIhbXw54Hk). For target capture baits used, see Medina et al. (2019) and Dryad Digital repository (https://datadryad.org/stash/dataset/doi:10.5061/dryad.8rq9465).

