## Supplementary Figures, Tables, and Files for "Frequent allopolyploidy with distant progenitors in the moss genera *Physcomitrium* and *Entosthodon* (Funariaceae) identified via subgenome phasing of targeted nuclear genes": SUPPLEMENTARY_FIGURES_ALL.pdf

Table S1. For each sample, collector and locality data, the total number of heterozygous sites, number of genes recovered, paralogy for all specimens.

I*psb*A-trnH and *rps5*-trnS.

Table S2. A list of gene names for the 50 nuclear loci analyzed using Homologizer, which were selected from the 648 gene set published in Medina et al. (2019).

Table S3. Primer sequences and citations for all sanger sequences markers for sample identity validation.

Table S4. Results from nQuire analysis of all samples. Values in columns 2-5 are likelihood values indicating fit of a free model and the lrdmodel for diploids, triploids, and tetraploids. Columns 6-8 indicate the delta-likelihood, or change in likelihood, between a free model and the lrdmodel for diploids, triploids, and tetraploids. Values in bold are the lowest likelihoods for each sample.

Figure S1. Phylogeny of the *Entosthodon-Physcomitrium* complex inferred from 50 genes and flanking regions. RevBayes tree including phased allopolyploid subgenomes sampled in the present study, with posterior probability indicated.

Figure S2. Phylogeny of the *Entosthodon-Physcomitrium* complex inferred from 648 nuclear genes and flanking regions. ASTRAL tree including all haploids and allopolyploids sampled, with all gene congruence indicated at each node. Pie charts on each branch indicate the proportion of gene trees concordant with that bipartition (blue), proportion of gene trees with the most common discordant bipartition for that bipartition (orange), proportion of gene trees with all other discordant bipartitions for that bipartition (red) and proportion of gene trees with no support due to missing data for that bipartition (black).

Figure S3: Alignment of chromatograms of single copy nuclear marker 7379 for *P. collenchymatum*-5274A and the associated voucher (*Homberg 1155*) and stock culture (IMSC 40061).

Figure S4. Phylogenetic reconstruction of haploid samples, phased diploid subgenomes, and three phased subgenomes of *P. pyriforme – 3410*, 3798 using ASTRAL from 50 gene trees, with allodiploids sungenome phases fixed (“iteration two“ analysis). Pie charts on each branch indicate the proportion of gene trees concordant with that bipartition (blue), proportion of gene trees with the most common discordant bipartition for that bipartition (orange), proportion of gene trees with all other discordant bipartitions for that bipartition (red) and proportion of gene trees with no support due to missing data for that bipartition (black). Boxes indicate named clades as discussed in text. Taxa in bold are allopolyploids represented by their maternal (M) and paternal (P) subgenomes.

Figure S5. Phylogeny of the *Entosthodon*-*Physcomitrium* complex inferred from 648 nuclear genes plus flanking regions. ASTRAL tree with branch lengths proportional to concordance.

Figure S6. Phylogeny of the *Entosthodon*-*Physcomitrium* complex inferred from 648 nuclear genes plus flanking regions. RevBayes tree with posterior probability support values. Allopolyploid subgenomes are highlighted with colors unique to individual allopolyploids.

Figure S7. Phylogeny of the *Entosthodon*-*Physcomitrium* complex inferred from 50 nuclear genes plus flanking regions with *P. pyriforme – 3410*, 3798 represented by three subgenome tips (“iteration two” analysis). ASTRAL tree with local posterior probability support values.

Figure S8. Posterior probability trace for Iteration three of homologizer.In this iteration, the phase for allodiploid samples were fixed to the subgenomes from iteration one. The putative allotriploids *P. pyriforme – 3410* and 3798 had three sequences per gene, plus a blank “dummy” sequence to allow for alternative locations of cryptic subgenomes. The trace summarizes posterior probability across 10 independent runs of RevBayeshomologizer, with 5000 generations each. There is a general lack of convergence both within runs and across runs.

Figure S9. Bayesian inference phylogeny based on plastid markers psbA-trnH and *rps*5-trnS. Sampling includes a subset of accessions included in Medina et al. (2019) and *P. collenchymatum*-5274 from the present study and is used to infer the maternal heritage of allopolyploid accessions of *P. collenchymatum* presented in Figure 3.

Figure S10. Phylogeny of the *Entosthodon-Physcomitrium* complex inferred from 648 nuclear genes and flanking regions. Astral tree including only haploids sampled in the present study, with all gene congruence indicated at each node. Pie charts on each branch indicate the proportion of gene trees concordant with that bipartition (blue), proportion of gene trees with the most common discordant bipartition for that bipartition (orange), proportion of gene trees with all other discordant bipartitions for that bipartition (red) and proportion of gene trees with no support due to missing data for that bipartition (black).

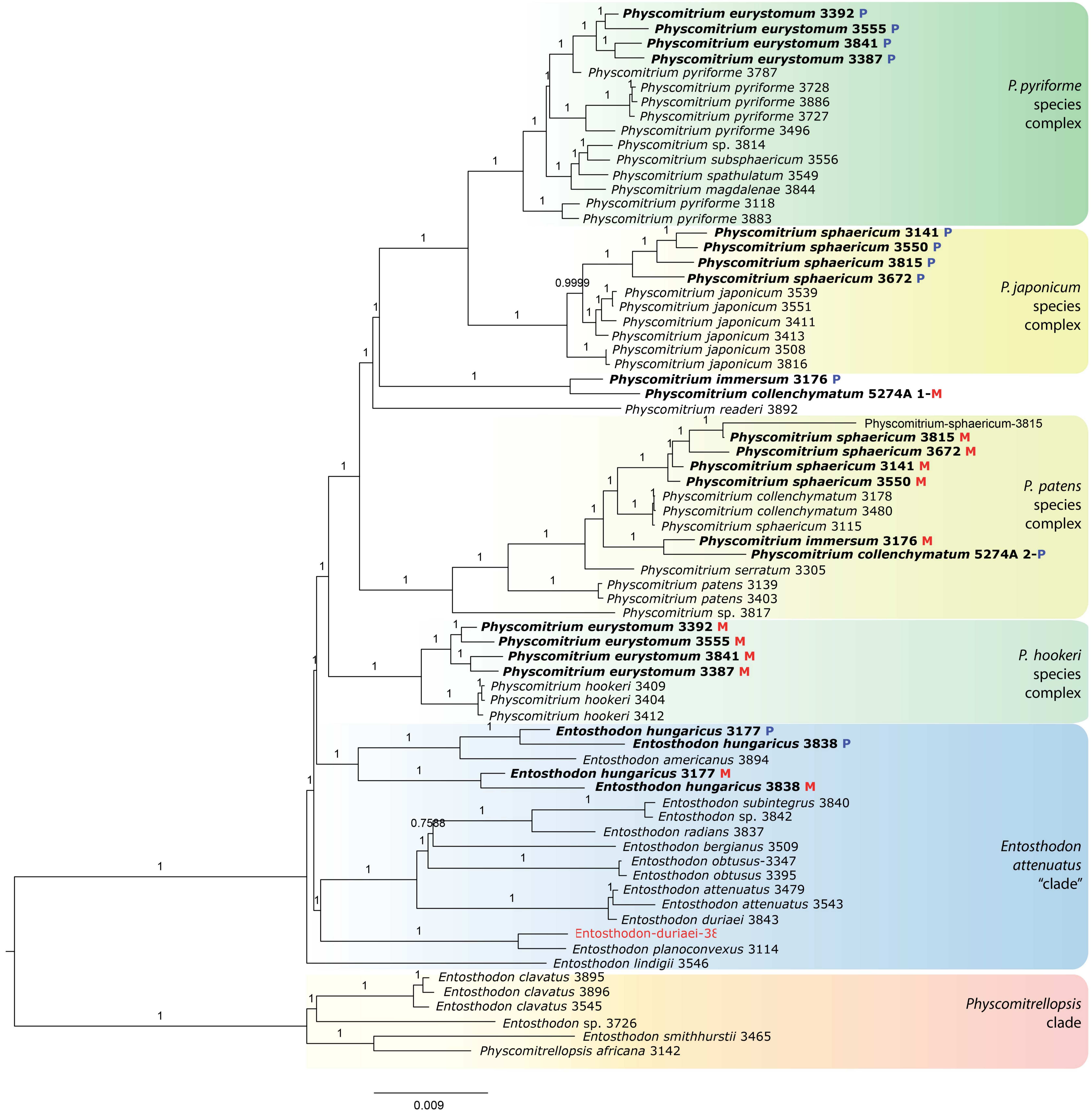

Physcomitrium eurystomum 3392 P
Physcomitrium eurystomum 3555 P
Physcomitrium eurystomum 3841 P
Physcomitrium eurystomum 3387 P
Physcomitrium pyriforme 3787
Physcomitrium pyriforme 3728
Physcomitrium pyriforme 3886
Physcomitrium pyriforme 3727
Physcomitrium pyriforme 3496
Physcomitrium sp. 3814
Physcomitrium subsphaericum 3556
Physcomitrium spathulatum 3549
Physcomitrium magdalenae 3844
Physcomitrium pyriforme 3118
Physcomitrium pyriforme 3883
P. pyriforme species complex
Physcomitrium sphaericum 3141 P
Physcomitrium sphaericum 3550 P
Physcomitrium sphaericum 3815 P
Physcomitrium sphaericum 3672 P
Physcomitrium japonicum 3539
Physcomitrium japonicum 3551
Physcomitrium japonicum 3411
Physcomitrium japonicum 3413
Physcomitrium japonicum 3508
Physcomitrium japonicum 3816
P. japonicum species complex
0.9999
Physcomitrium immersum 3176 P
Physcomitrium collenchymatum 5274A 1-M
Physcomitrium readeri 3892
Physcomitrium-sphaericum-3815
Physcomitrium sphaericum 3815 M
Physcomitrium sphaericum 3672 M
Physcomitrium sphaericum 3141 M
Physcomitrium sphaericum 3550 M
Physcomitrium collenchymatum 3178
Physcomitrium collenchymatum 3480
Physcomitrium sphaericum 3115
Physcomitrium immersum 3176 M
Physcomitrium collenchymatum 5274A 2-P
Physcomitrium serratum 3305
Physcomitrium patens 3139
Physcomitrium patens 3403
Physcomitrium sp. 3817
P. patens species complex
Physcomitrium eurystomum 3392 M
Physcomitrium eurystomum 3555 M
Physcomitrium eurystomum 3841 M
Physcomitrium eurystomum 3387 M
Physcomitrium hookeri 3409
Physcomitrium hookeri 3404
Physcomitrium hookeri 3412
P. hookeri species complex
Entosthodon hungaricus 3177 P
Entosthodon hungaricus 3838 P
Entosthodon americanus 3894
Entosthodon hungaricus 3177 M
Entosthodon hungaricus 3838 M
Entosthodon subintegrus 3840
Entosthodon sp. 3842
Entosthodon radians 3837
0.7588
Entosthodon bergianus 3509
Entosthodon obtusus-3347
Entosthodon obtusus 3395
Entosthodon attenuatus 3479
Entosthodon attenuatus 3543
Entosthodon duriaei 3843
Entosthodon-duriaei-3ε
Entosthodon planoconvexus 3114
Entosthodon lindigii 3546
Entosthodon attenuatus "clade"
Entosthodon clavatus 3895
Entosthodon clavatus 3896
Entosthodon clavatus 3545
Entosthodon sp. 3726
Entosthodon smithhurstii 3465
Physcomitrellopsis africana 3142
Physcomitrellopsis clade
0.009

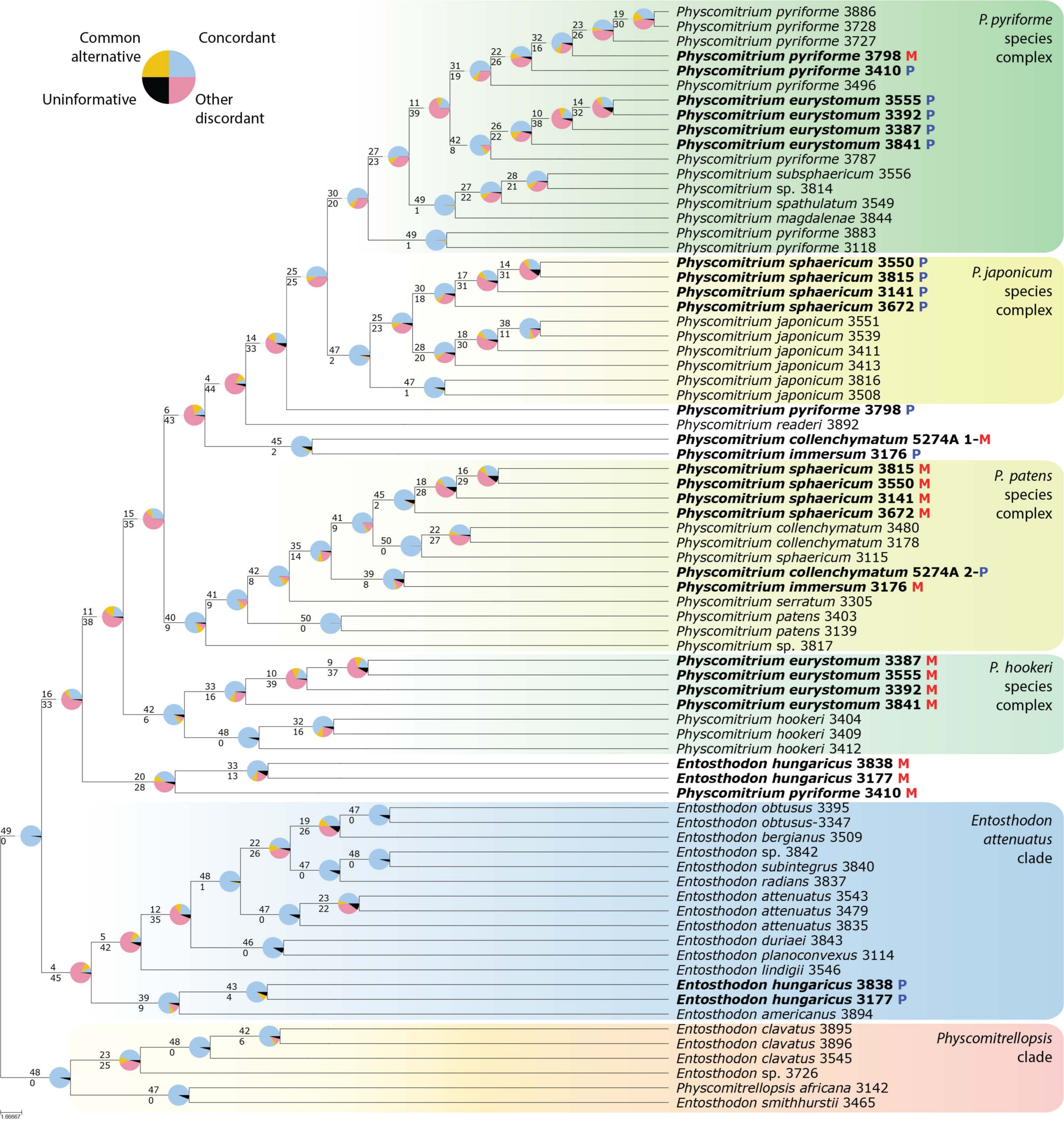
Common alternative
Concordant
Uninformative
Other discordant
P. pyriforme species complex
P. japonicum species complex
P. patens species complex
P. hookeri species complex
Entosthodon attenuatus clade
Physcomitrellopsis clade
Physcomitrium pyriforme 3886
Physcomitrium pyriforme 3728
Physcomitrium pyriforme 3727
Physcomitrium pyriforme 3798 M
Physcomitrium pyriforme 3410 P
Physcomitrium pyriforme 3496
Physcomitrium eurystomum 3555 P
Physcomitrium eurystomum 3392 P
Physcomitrium eurystomum 3387 P
Physcomitrium eurystomum 3841 P
Physcomitrium pyriforme 3787
Physcomitrium subsphaericum 3556
Physcomitrium sp. 3814
Physcomitrium spathulatum 3549
Physcomitrium magdalenae 3844
Physcomitrium pyriforme 3883
Physcomitrium pyriforme 3118
Physcomitrium sphaericum 3550 P
Physcomitrium sphaericum 3815 P
Physcomitrium sphaericum 3141 P
Physcomitrium sphaericum 3672 P
Physcomitrium japonicum 3551
Physcomitrium japonicum 3539
Physcomitrium japonicum 3411
Physcomitrium japonicum 3413
Physcomitrium japonicum 3816
Physcomitrium japonicum 3508
Physcomitrium pyriforme 3798 P
Physcomitrium readeri 3892
Physcomitrium collenchymatum 5274A 1-M
Physcomitrium immersum 3176 P
Physcomitrium sphaericum 3815 M
Physcomitrium sphaericum 3550 M
Physcomitrium sphaericum 3141 M
Physcomitrium sphaericum 3672 M
Physcomitrium collenchymatum 3480
Physcomitrium collenchymatum 3178
Physcomitrium sphaericum 3115
Physcomitrium collenchymatum 5274A 2-P
Physcomitrium immersum 3176 M
Physcomitrium serratum 3305
Physcomitrium patens 3403
Physcomitrium patens 3139
Physcomitrium sp. 3817
Physcomitrium eurystomum 3387 M
Physcomitrium eurystomum 3555 M
Physcomitrium eurystomum 3392 M
Physcomitrium eurystomum 3841 M
Physcomitrium hookeri 3404
Physcomitrium hookeri 3409
Physcomitrium hookeri 3412
Entosthodon hungaricus 3838 M
Entosthodon hungaricus 3177 M
Physcomitrium pyriforme 3410 M
Entosthodon obtusus 3395
Entosthodon obtusus-3347
Entosthodon bergianus 3509
Entosthodon sp. 3842
Entosthodon subintegrus 3840
Entosthodon radians 3837
Entosthodon attenuatus 3543
Entosthodon attenuatus 3479
Entosthodon attenuatus 3835
Entosthodon duriaei 3843
Entosthodon planoconvexus 3114
Entosthodon lindigii 3546
Entosthodon hungaricus 3838 P
Entosthodon hungaricus 3177 P
Entosthodon americanus 3894
Entosthodon clavatus 3895
Entosthodon clavatus 3896
Entosthodon clavatus 3545
Entosthodon sp. 3726
Physcomitrellopsis africana 3142
Entosthodon smithhurstii 3465
1.66667

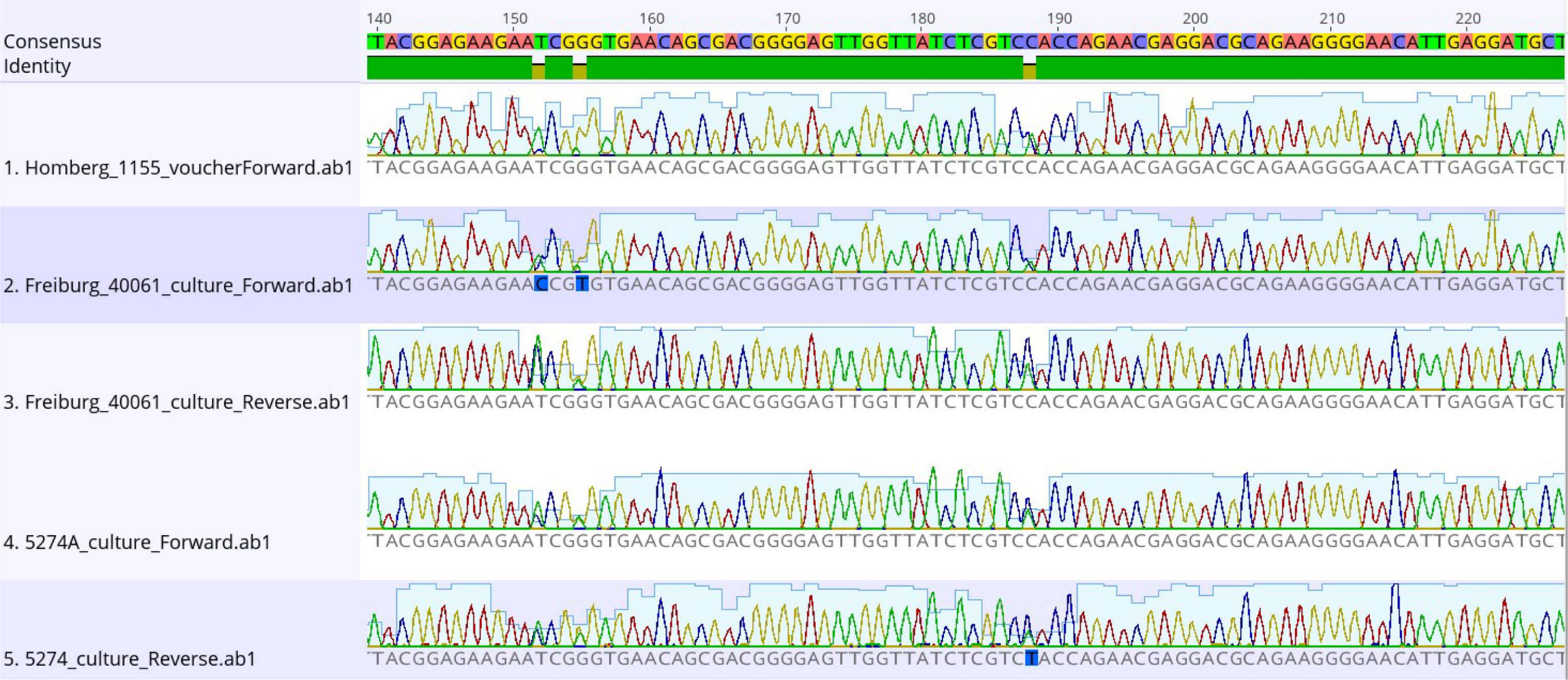

140
150
160
170
180
190
200
210
220
Consensus
TACGGAGAAGAATCGGGTGAACAGCGACGGGGAGTTGGTTATCTCGTCCACCAGAACGAGGACGCAGAAGGGGAACATTGAGGATGCT
Identity
1. Homberg_1155_voucherForward.ab1
TACGGAGAAGAATCGGGTGAACAGCGACGGGGAGTTGGTTATCTCGTCCACCAGAACGAGGACGCAGAAGGGGAACATTGAGGATGCT
2. Freiburg_40061_culture_Forward.ab1
TACGGAGAAGAACCGTGTGAACAGCGACGGGGAGTTGGTTATCTCGTCCACCAGAACGAGGACGCAGAAGGGGAACATTGAGGATGCT
3. Freiburg_40061_culture_Reverse.ab1
TACGGAGAAGAATCGGGTGAACAGCGACGGGGAGTTGGTTATCTCGTCCACCAGAACGAGGACGCAGAAGGGGAACATTGAGGATGCT
4. 5274A_culture_Forward.ab1
TACGGAGAAGAATCGGGTGAACAGCGACGGGGAGTTGGTTATCTCGTCCACCAGAACGAGGACGCAGAAGGGGAACATTGAGGATGCT
5. 5274_culture_Reverse.ab1
TACGGAGAAGAATCGGGTGAACAGCGACGGGGAGTTGGTTATCTCGTCTACCAGAACGAGGACGCAGAAGGGGAACATTGAGGATGCT

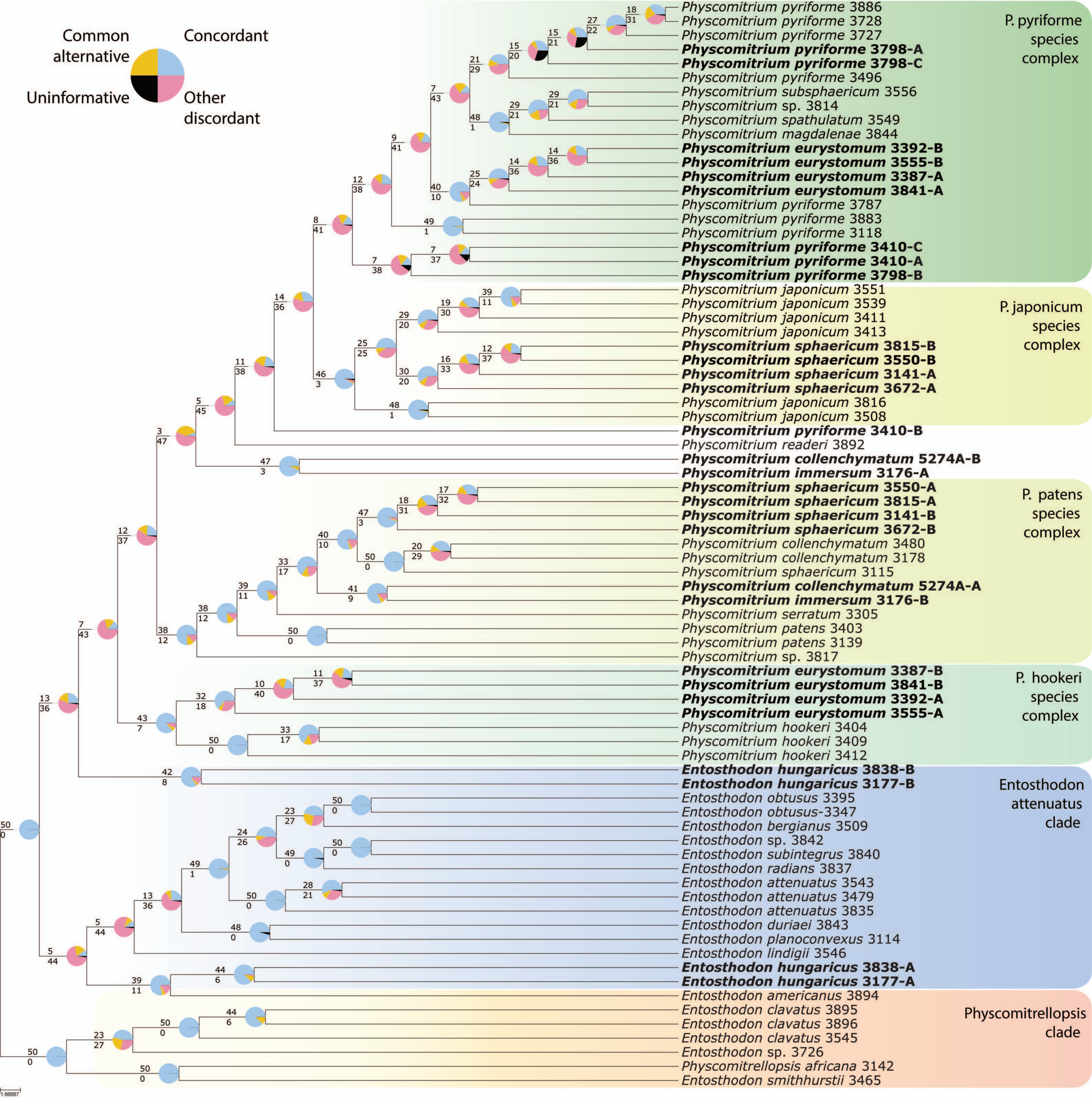

Common alternative
Concordant
Uninformative
Other discordant
P. pyriforme species complex
P. japonicum species complex
P. patens species complex
P. hookeri species complex
Entosthodon attenuatus clade
Physcomitrellopsis clade
Physcomitrium pyriforme 3886
Physcomitrium pyriforme 3728
Physcomitrium pyriforme 3727
Physcomitrium pyriforme 3798-A
Physcomitrium pyriforme 3798-C
Physcomitrium pyriforme 3496
Physcomitrium subsphaericum 3556
Physcomitrium sp. 3814
Physcomitrium spathulatum 3549
Physcomitrium magdalenae 3844
Physcomitrium eurystomum 3392-B
Physcomitrium eurystomum 3555-B
Physcomitrium eurystomum 3387-A
Physcomitrium eurystomum 3841-A
Physcomitrium pyriforme 3787
Physcomitrium pyriforme 3883
Physcomitrium pyriforme 3118
Physcomitrium pyriforme 3410-C
Physcomitrium pyriforme 3410-A
Physcomitrium pyriforme 3798-B
Physcomitrium japonicum 3551
Physcomitrium japonicum 3539
Physcomitrium japonicum 3411
Physcomitrium japonicum 3413
Physcomitrium sphaericum 3815-B
Physcomitrium sphaericum 3550-B
Physcomitrium sphaericum 3141-A
Physcomitrium sphaericum 3672-A
Physcomitrium japonicum 3816
Physcomitrium japonicum 3508
Physcomitrium pyriforme 3410-B
Physcomitrium readeri 3892
Physcomitrium collenchymatum 5274A-B
Physcomitrium immersum 3176-A
Physcomitrium sphaericum 3550-A
Physcomitrium sphaericum 3815-A
Physcomitrium sphaericum 3141-B
Physcomitrium sphaericum 3672-B
Physcomitrium collenchymatum 3480
Physcomitrium collenchymatum 3178
Physcomitrium sphaericum 3115
Physcomitrium collenchymatum 5274A-A
Physcomitrium immersum 3176-B
Physcomitrium serratum 3305
Physcomitrium patens 3403
Physcomitrium patens 3139
Physcomitrium sp. 3817
Physcomitrium eurystomum 3387-B
Physcomitrium eurystomum 3841-B
Physcomitrium eurystomum 3392-A
Physcomitrium eurystomum 3555-A
Physcomitrium hookeri 3404
Physcomitrium hookeri 3409
Physcomitrium hookeri 3412
Entosthodon hungaricus 3838-B
Entosthodon hungaricus 3177-B
Entosthodon obtusus 3395
Entosthodon obtusus-3347
Entosthodon bergianus 3509
Entosthodon sp. 3842
Entosthodon subintegrus 3840
Entosthodon radians 3837
Entosthodon attenuatus 3543
Entosthodon attenuatus 3479
Entosthodon attenuatus 3835
Entosthodon duriaei 3843
Entosthodon planoconvexus 3114
Entosthodon lindigii 3546
Entosthodon hungaricus 3838-A
Entosthodon hungaricus 3177-A
Entosthodon americanus 3894
Entosthodon clavatus 3895
Entosthodon clavatus 3896
Entosthodon clavatus 3545
Entosthodon sp. 3726
Physcomitrellopsis africana 3142
Entosthodon smithhurstii 3465
1.66667

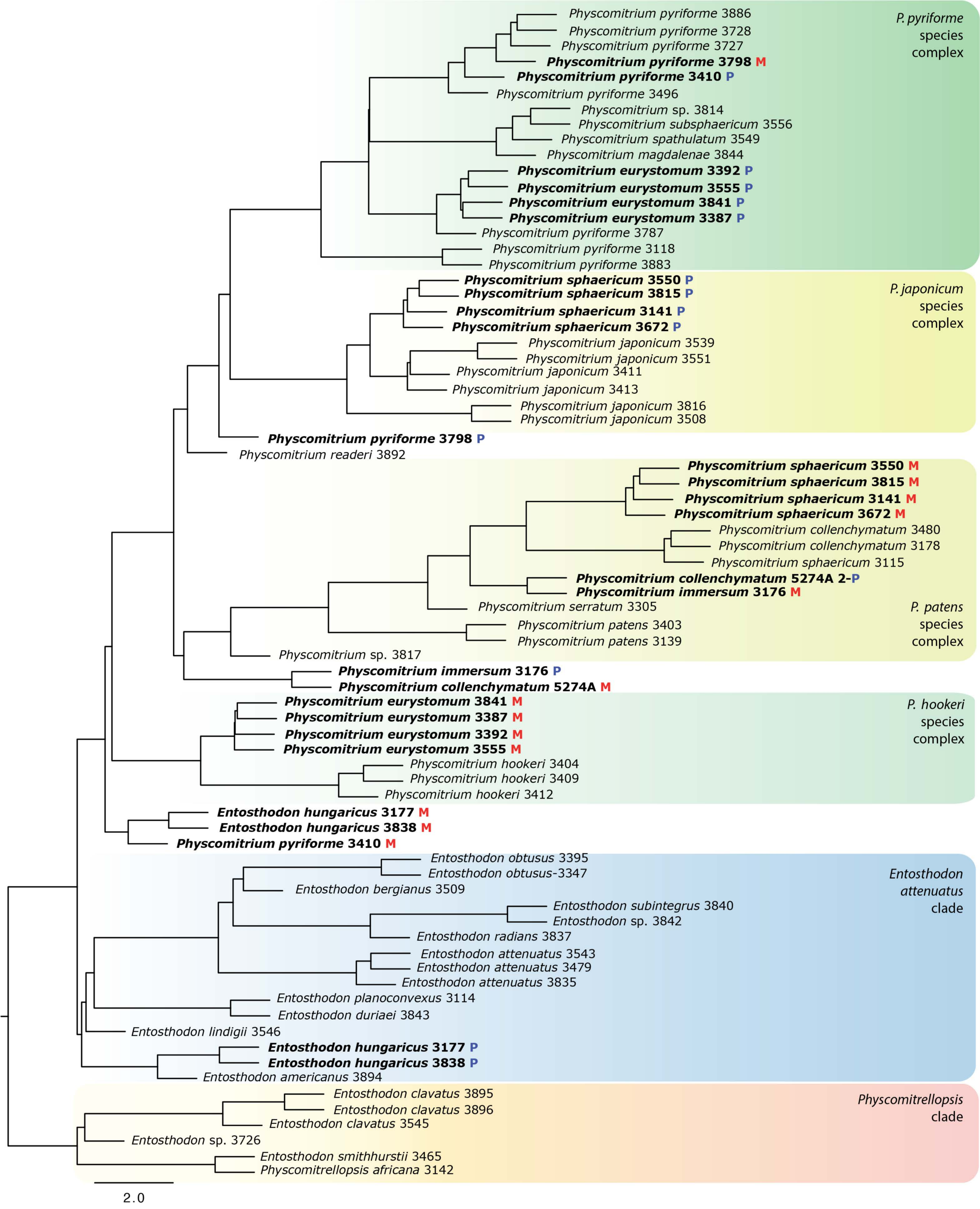

Physcomitrium pyriforme 3886
Physcomitrium pyriforme 3728
Physcomitrium pyriforme 3727
Physcomitrium pyriforme 3798 M
Physcomitrium pyriforme 3410 P
Physcomitrium pyriforme 3496
Physcomitrium sp. 3814
Physcomitrium subsphaericum 3556
Physcomitrium spathulatum 3549
Physcomitrium magdalenae 3844
Physcomitrium eurystomum 3392 P
Physcomitrium eurystomum 3555 P
Physcomitrium eurystomum 3841 P
Physcomitrium eurystomum 3387 P
Physcomitrium pyriforme 3787
Physcomitrium pyriforme 3118
Physcomitrium pyriforme 3883
P. pyriforme species complex
Physcomitrium sphaericum 3550 P
Physcomitrium sphaericum 3815 P
Physcomitrium sphaericum 3141 P
Physcomitrium sphaericum 3672 P
Physcomitrium japonicum 3539
Physcomitrium japonicum 3551
Physcomitrium japonicum 3411
Physcomitrium japonicum 3413
Physcomitrium japonicum 3816
Physcomitrium japonicum 3508
P. japonicum species complex
Physcomitrium pyriforme 3798 P
Physcomitrium readeri 3892
Physcomitrium sphaericum 3550 M
Physcomitrium sphaericum 3815 M
Physcomitrium sphaericum 3141 M
Physcomitrium sphaericum 3672 M
Physcomitrium collenchymatum 3480
Physcomitrium collenchymatum 3178
Physcomitrium sphaericum 3115
Physcomitrium collenchymatum 5274A 2-P
Physcomitrium immersum 3176 M
Physcomitrium serratum 3305
Physcomitrium patens 3403
Physcomitrium patens 3139
Physcomitrium sp. 3817
P. patens species complex
Physcomitrium immersum 3176 P
Physcomitrium collenchymatum 5274A M
Physcomitrium eurystomum 3841 M
Physcomitrium eurystomum 3387 M
Physcomitrium eurystomum 3392 M
Physcomitrium eurystomum 3555 M
Physcomitrium hookeri 3404
Physcomitrium hookeri 3409
Physcomitrium hookeri 3412
P. hookeri species complex
Entosthodon hungaricus 3177 M
Entosthodon hungaricus 3838 M
Physcomitrium pyriforme 3410 M
Entosthodon obtusus 3395
Entosthodon obtusus-3347
Entosthodon bergianus 3509
Entosthodon subintegrus 3840
Entosthodon sp. 3842
Entosthodon radians 3837
Entosthodon attenuatus 3543
Entosthodon attenuatus 3479
Entosthodon attenuatus 3835
Entosthodon planoconvexus 3114
Entosthodon duriaei 3843
Entosthodon lindigii 3546
Entosthodon hungaricus 3177 P
Entosthodon hungaricus 3838 P
Entosthodon americanus 3894
Entosthodon attenuatus clade
Entosthodon clavatus 3895
Entosthodon clavatus 3896
Entosthodon clavatus 3545
Entosthodon sp. 3726
Entosthodon smithhurstii 3465
Physcomitrellopsis africana 3142
Physcomitrellopsis clade
2.0

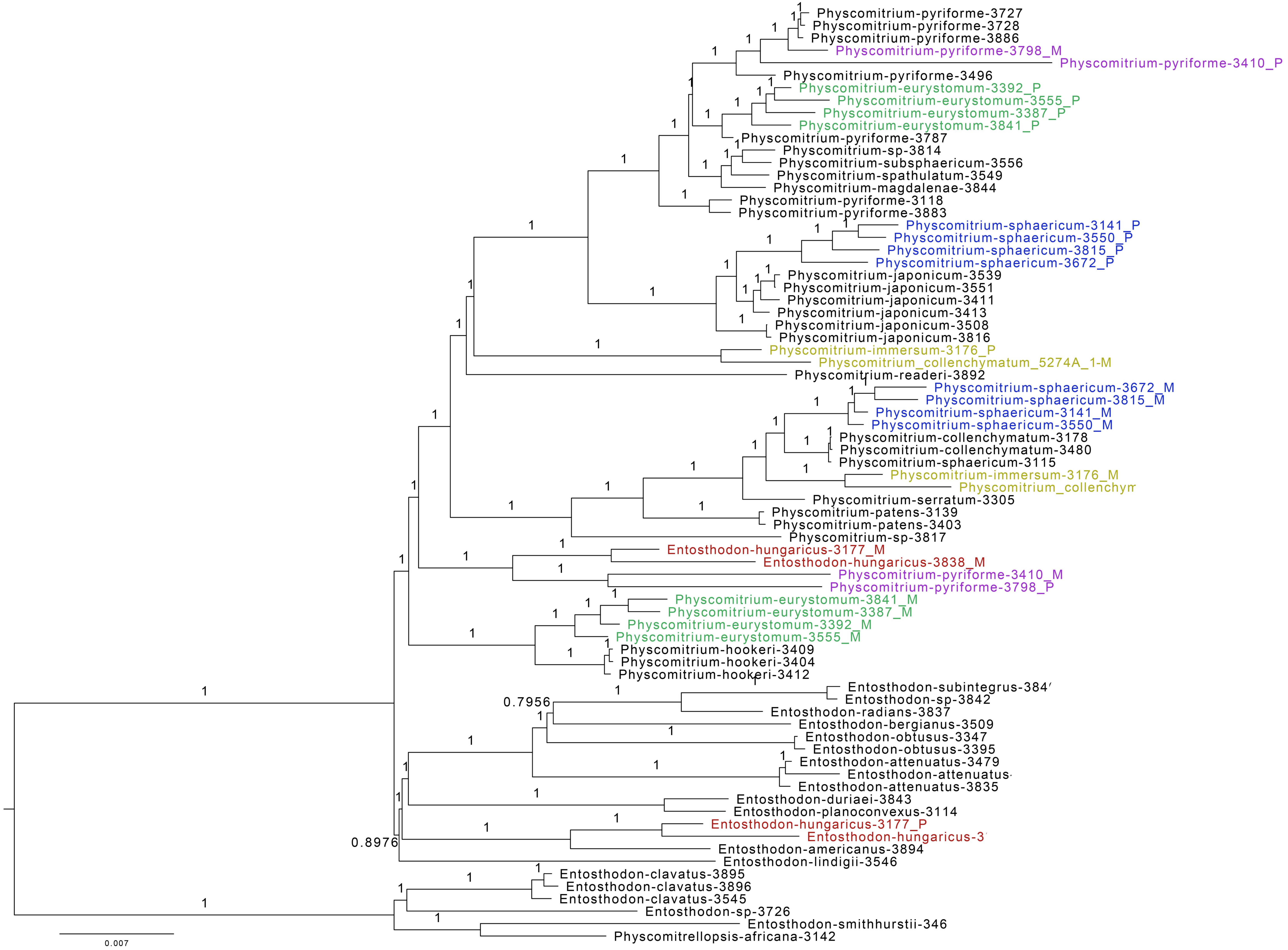

Physcomitrium-pyriforme-3727
Physcomitrium-pyriforme-3728
Physcomitrium-pyriforme-3886
Physcomitrium-pyriforme-3798_M
Physcomitrium-pyriforme-3410_P
Physcomitrium-pyriforme-3496
Physcomitrium-eurystomum-3392_P
Physcomitrium-eurystomum-3555_P
Physcomitrium-eurystomum-3387_P
Physcomitrium-eurystomum-3841_P
Physcomitrium-pyriforme-3787
Physcomitrium-sp-3814
Physcomitrium-subsphaericum-3556
Physcomitrium-spathulatum-3549
Physcomitrium-magdalenae-3844
Physcomitrium-pyriforme-3118
Physcomitrium-pyriforme-3883
Physcomitrium-sphaericum-3141_P
Physcomitrium-sphaericum-3550_P
Physcomitrium-sphaericum-3815_P
Physcomitrium-sphaericum-3672_P
Physcomitrium-japonicum-3539
Physcomitrium-japonicum-3551
Physcomitrium-japonicum-3411
Physcomitrium-japonicum-3413
Physcomitrium-japonicum-3508
Physcomitrium-japonicum-3816
Physcomitrium-immersum-3176_P
Physcomitrium_collenchymatum_5274A_1-M
Physcomitrium-readeri-3892
Physcomitrium-sphaericum-3672_M
Physcomitrium-sphaericum-3815_M
Physcomitrium-sphaericum-3141_M
Physcomitrium-sphaericum-3550_M
Physcomitrium-collenchymatum-3178
Physcomitrium-collenchymatum-3480
Physcomitrium-sphaericum-3115
Physcomitrium-immersum-3176_M
Physcomitrium_collenchym
Physcomitrium-serratum-3305
Physcomitrium-patens-3139
Physcomitrium-patens-3403
Physcomitrium-sp-3817
Entosthodon-hungaricus-3177_M
Entosthodon-hungaricus-3838_M
Physcomitrium-pyriforme-3410_M
Physcomitrium-pyriforme-3798_P
Physcomitrium-eurystomum-3841_M
Physcomitrium-eurystomum-3387_M
Physcomitrium-eurystomum-3392_M
Physcomitrium-eurystomum-3555_M
Physcomitrium-hookeri-3409
Physcomitrium-hookeri-3404
Physcomitrium-hookeri-3412
Entosthodon-subintegrus-384
Entosthodon-sp-3842
Entosthodon-radians-3837
Entosthodon-bergianus-3509
Entosthodon-obtusus-3347
Entosthodon-obtusus-3395
Entosthodon-attenuatus-3479
Entosthodon-attenuatus
Entosthodon-attenuatus-3835
Entosthodon-duriaei-3843
Entosthodon-planoconvexus-3114
Entosthodon-hungaricus-3177_P
Entosthodon-hungaricus-3
Entosthodon-americanus-3894
Entosthodon-lindigii-3546
Entosthodon-clavatus-3895
Entosthodon-clavatus-3896
Entosthodon-clavatus-3545
Entosthodon-sp-3726
Entosthodon-smithhurstii-346
Physcomitrellopsis-africana-3142
0.7956
0.8976
0.007

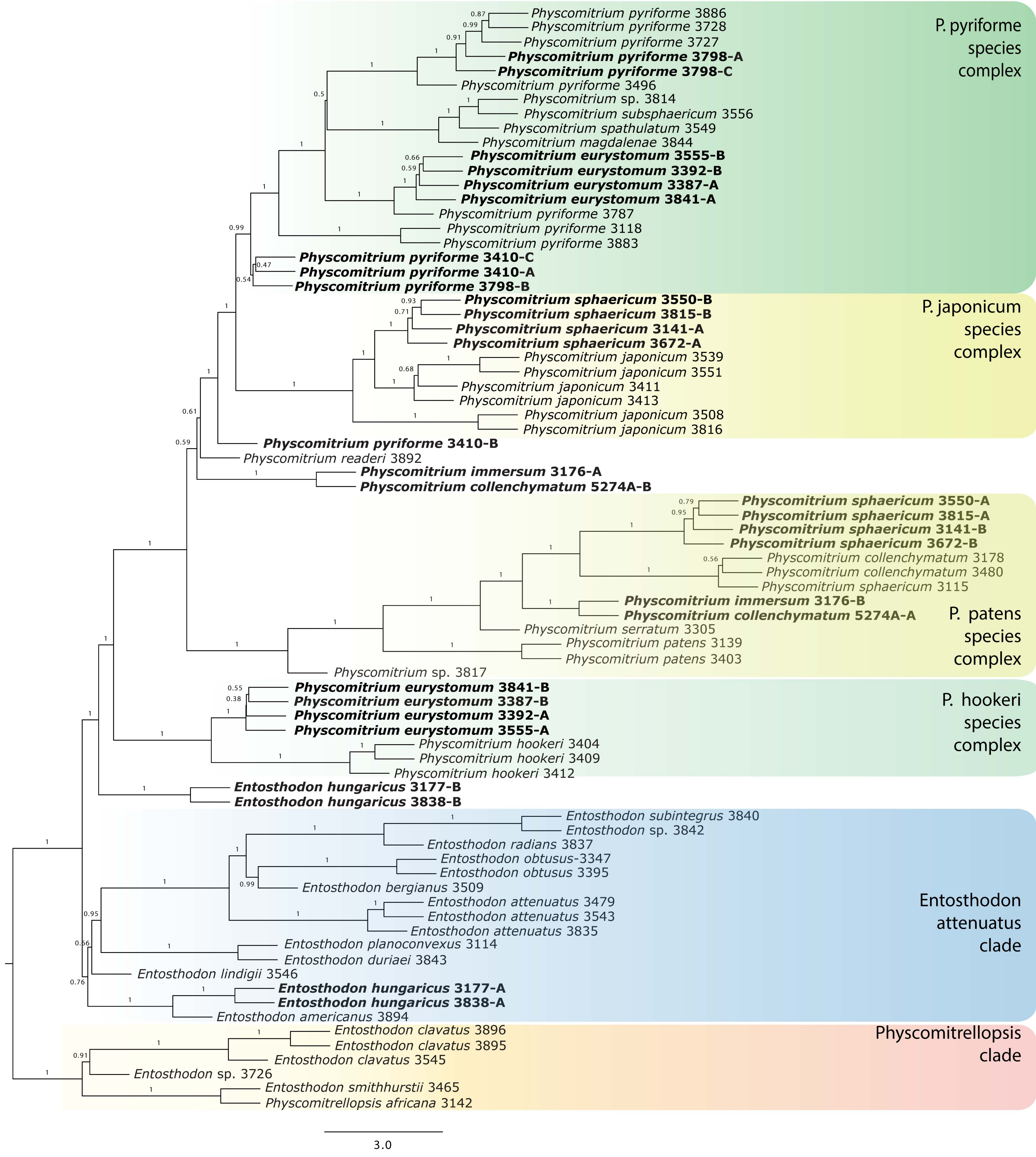

Physcomitrium pyriforme 3886
Physcomitrium pyriforme 3728
Physcomitrium pyriforme 3727
Physcomitrium pyriforme 3798-A
Physcomitrium pyriforme 3798-C
Physcomitrium pyriforme 3496
Physcomitrium sp. 3814
Physcomitrium subsphaericum 3556
Physcomitrium spathulatum 3549
Physcomitrium magdalenae 3844
Physcomitrium eurystomum 3555-B
Physcomitrium eurystomum 3392-B
Physcomitrium eurystomum 3387-A
Physcomitrium eurystomum 3841-A
Physcomitrium pyriforme 3787
Physcomitrium pyriforme 3118
Physcomitrium pyriforme 3883
Physcomitrium pyriforme 3410-C
Physcomitrium pyriforme 3410-A
Physcomitrium pyriforme 3798-B
P. pyriforme species complex
Physcomitrium sphaericum 3550-B
Physcomitrium sphaericum 3815-B
Physcomitrium sphaericum 3141-A
Physcomitrium sphaericum 3672-A
Physcomitrium japonicum 3539
Physcomitrium japonicum 3551
Physcomitrium japonicum 3411
Physcomitrium japonicum 3413
Physcomitrium japonicum 3508
Physcomitrium japonicum 3816
P. japonicum species complex
Physcomitrium pyriforme 3410-B
Physcomitrium readeri 3892
Physcomitrium immersum 3176-A
Physcomitrium collenchymatum 5274A-B
Physcomitrium sphaericum 3550-A
Physcomitrium sphaericum 3815-A
Physcomitrium sphaericum 3141-B
Physcomitrium sphaericum 3672-B
Physcomitrium collenchymatum 3178
Physcomitrium collenchymatum 3480
Physcomitrium sphaericum 3115
Physcomitrium immersum 3176-B
Physcomitrium collenchymatum 5274A-A
Physcomitrium serratum 3305
Physcomitrium patens 3139
Physcomitrium patens 3403
Physcomitrium sp. 3817
P. patens species complex
Physcomitrium eurystomum 3841-B
Physcomitrium eurystomum 3387-B
Physcomitrium eurystomum 3392-A
Physcomitrium eurystomum 3555-A
Physcomitrium hookeri 3404
Physcomitrium hookeri 3409
Physcomitrium hookeri 3412
P. hookeri species complex
Entosthodon hungaricus 3177-B
Entosthodon hungaricus 3838-B
Entosthodon subintegrus 3840
Entosthodon sp. 3842
Entosthodon radians 3837
Entosthodon obtusus-3347
Entosthodon obtusus 3395
Entosthodon bergianus 3509
Entosthodon attenuatus 3479
Entosthodon attenuatus 3543
Entosthodon attenuatus 3835
Entosthodon planoconvexus 3114
Entosthodon duriaei 3843
Entosthodon lindigii 3546
Entosthodon hungaricus 3177-A
Entosthodon hungaricus 3838-A
Entosthodon americanus 3894
Entosthodon attenuatus clade
Entosthodon clavatus 3896
Entosthodon clavatus 3895
Entosthodon clavatus 3545
Entosthodon sp. 3726
Entosthodon smithhurstii 3465
Physcomitrellopsis africana 3142
Physcomitrellopsis clade
3.0

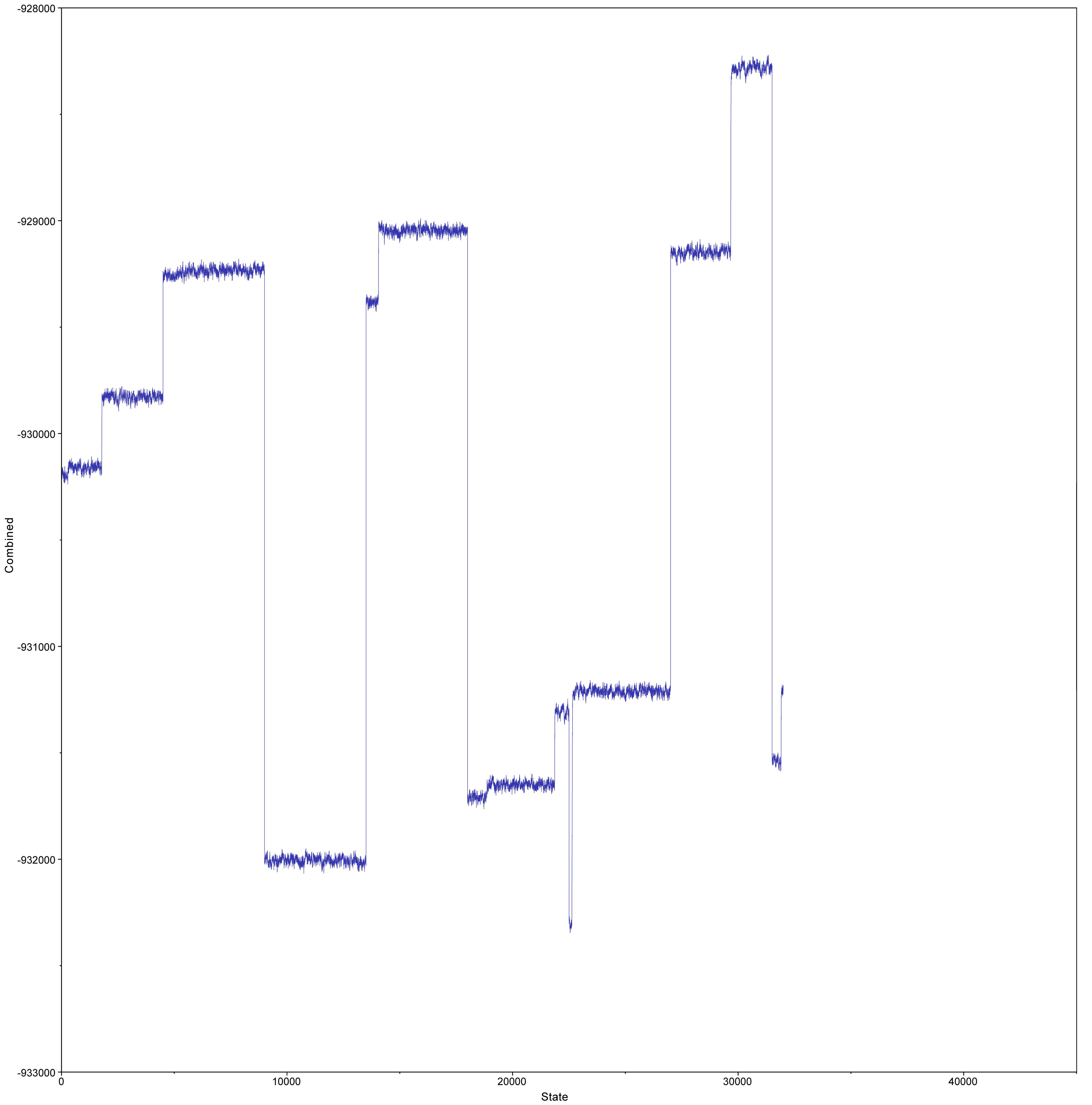

-928000
-929000
-930000
-931000
-932000
-933000
Combined
0
10000
20000
30000
40000
State

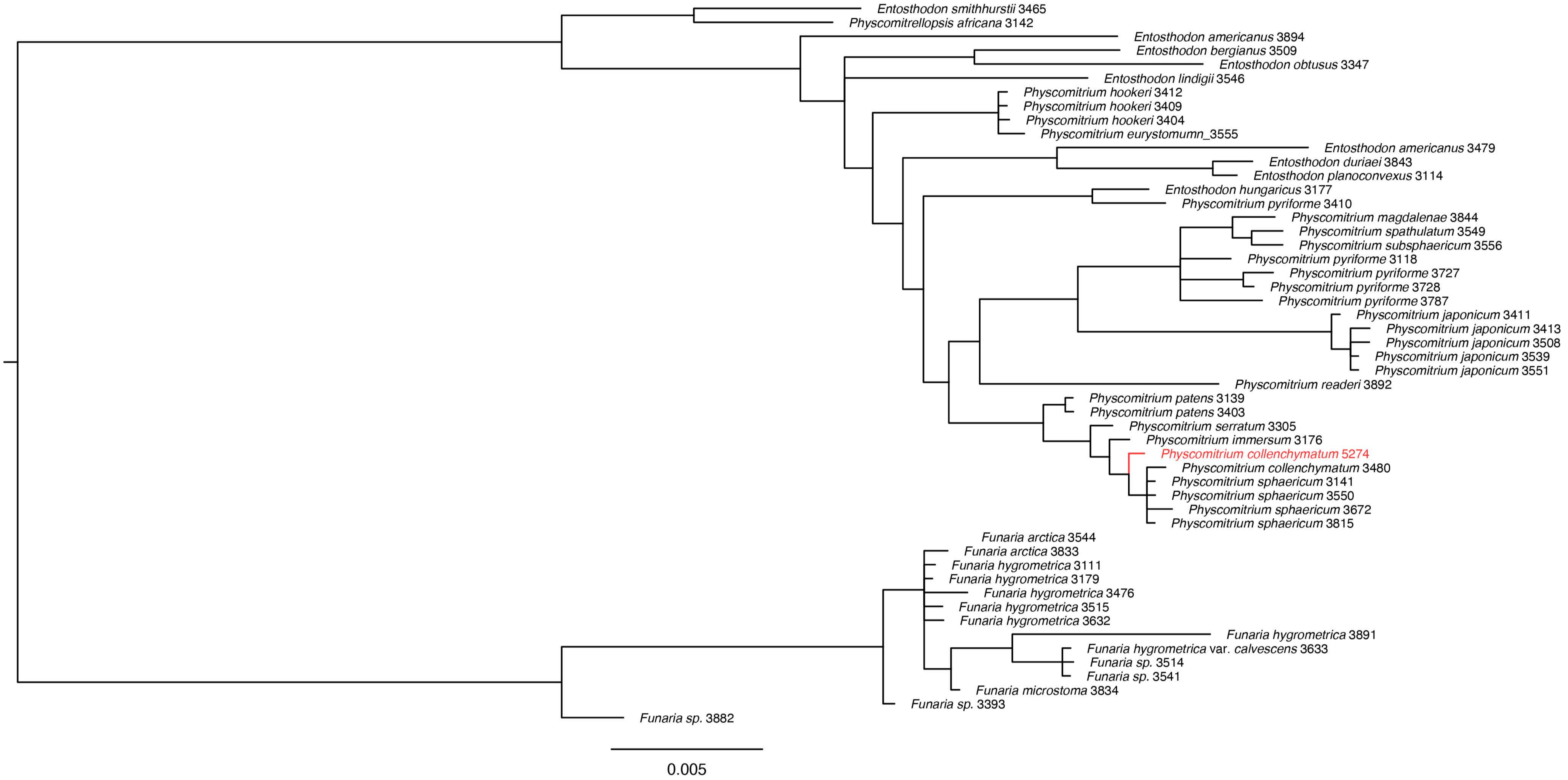

Entosthodon smithhurstii 3465
Physcomitrellopsis africana 3142
Entosthodon americanus 3894
Entosthodon bergianus 3509
Entosthodon obtusus 3347
Entosthodon lindigii 3546
Physcomitrium hookeri 3412
Physcomitrium hookeri 3409
Physcomitrium hookeri 3404
Physcomitrium eurystomumn_3555
Entosthodon americanus 3479
Entosthodon duriaei 3843
Entosthodon planoconvexus 3114
Entosthodon hungaricus 3177
Physcomitrium pyriforme 3410
Physcomitrium magdalenae 3844
Physcomitrium spathulatum 3549
Physcomitrium subsphaericum 3556
Physcomitrium pyriforme 3118
Physcomitrium pyriforme 3727
Physcomitrium pyriforme 3728
Physcomitrium pyriforme 3787
Physcomitrium japonicum 3411
Physcomitrium japonicum 3413
Physcomitrium japonicum 3508
Physcomitrium japonicum 3539
Physcomitrium japonicum 3551
Physcomitrium readeri 3892
Physcomitrium patens 3139
Physcomitrium patens 3403
Physcomitrium serratum 3305
Physcomitrium immersum 3176
Physcomitrium collenchymatum 5274
Physcomitrium collenchymatum 3480
Physcomitrium sphaericum 3141
Physcomitrium sphaericum 3550
Physcomitrium sphaericum 3672
Physcomitrium sphaericum 3815
Funaria arctica 3544
Funaria arctica 3833
Funaria hygrometrica 3111
Funaria hygrometrica 3179
Funaria hygrometrica 3476
Funaria hygrometrica 3515
Funaria hygrometrica 3632
Funaria hygrometrica 3891
Funaria hygrometrica var. calvescens 3633
Funaria sp. 3514
Funaria sp. 3541
Funaria microstoma 3834
Funaria sp. 3393
Funaria sp. 3882
0.005

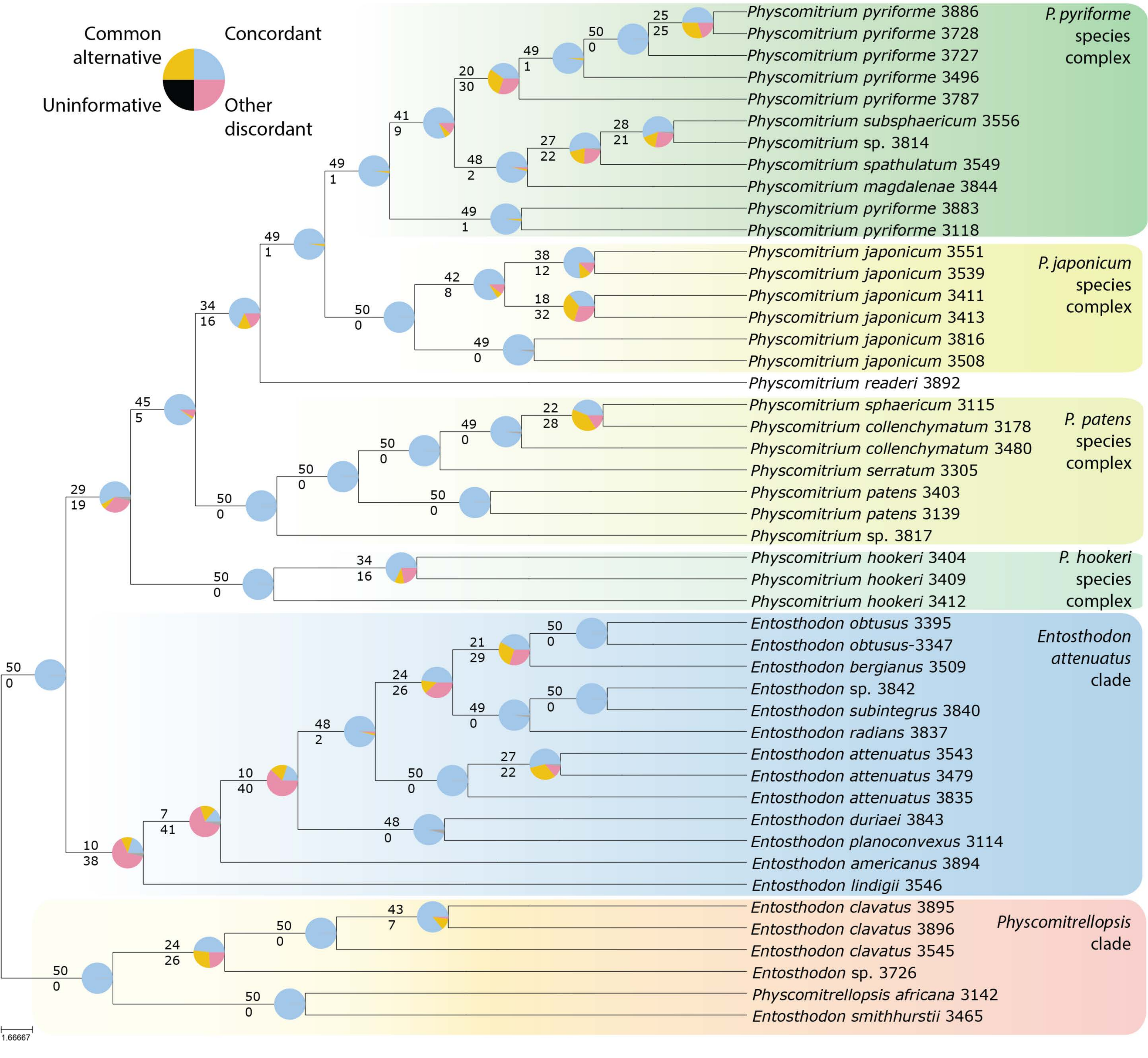
Common alternative
Concordant
Uninformative
Other discordant
Physcomitrium pyriforme 3886
Physcomitrium pyriforme 3728
Physcomitrium pyriforme 3727
Physcomitrium pyriforme 3496
Physcomitrium pyriforme 3787
Physcomitrium subsphaericum 3556
Physcomitrium sp. 3814
Physcomitrium spathulatum 3549
Physcomitrium magdalenae 3844
Physcomitrium pyriforme 3883
Physcomitrium pyriforme 3118
P. pyriforme species complex
Physcomitrium japonicum 3551
Physcomitrium japonicum 3539
Physcomitrium japonicum 3411
Physcomitrium japonicum 3413
Physcomitrium japonicum 3816
Physcomitrium japonicum 3508
P. japonicum species complex
Physcomitrium readeri 3892
Physcomitrium sphaericum 3115
Physcomitrium collenchymatum 3178
Physcomitrium collenchymatum 3480
Physcomitrium serratum 3305
Physcomitrium patens 3403
Physcomitrium patens 3139
Physcomitrium sp. 3817
P. patens species complex
Physcomitrium hookeri 3404
Physcomitrium hookeri 3409
Physcomitrium hookeri 3412
P. hookeri species complex
Entosthodon obtusus 3395
Entosthodon obtusus-3347
Entosthodon bergianus 3509
Entosthodon sp. 3842
Entosthodon subintegrus 3840
Entosthodon radians 3837
Entosthodon attenuatus 3543
Entosthodon attenuatus 3479
Entosthodon attenuatus 3835
Entosthodon duriaei 3843
Entosthodon planoconvexus 3114
Entosthodon americanus 3894
Entosthodon lindigii 3546
Entosthodon attenuatus clade
Entosthodon clavatus 3895
Entosthodon clavatus 3896
Entosthodon clavatus 3545
Entosthodon sp. 3726
Physcomitrellopsis africana 3142
Entosthodon smithhurstii 3465
Physcomitrellopsis clade
1.66667
